# Developmental origins of Parkinson’s disease risk: perinatal exposure to the organochlorine pesticide dieldrin leads to sex-specific DNA modifications in critical neurodevelopmental pathways in the mouse midbrain

**DOI:** 10.1101/2024.04.26.590998

**Authors:** Joseph Kochmanski, Mahek Virani, Nathan C. Kuhn, Sierra L. Boyd, Katelyn Becker, Marie Adams, Alison I. Bernstein

**Author notes:** **Corresponding Author:** Alison Bernstein, Environmental and Occupational Health Sciences Institute, Ernest Mario School of Pharmacy, Rutgers University 170 Freylinghuysen Rd, Piscataway, NJ 08854.

## Abstract

Epidemiological studies show that exposure to the organochlorine pesticide dieldrin is associated with increased risk of Parkinson’s disease (PD). Animal studies support a link between developmental dieldrin exposure and increased neuronal susceptibility in the α-synuclein preformed fibril (α-syn PFF) and MPTP models in adult male C57BL/6 mice. In a previous study, we showed that developmental dieldrin exposure was associated with sex-specific changes in DNA modifications within genes related to dopaminergic neuron development and maintenance at 12 weeks of age. Here, we used capture hybridization-sequencing with custom baits to interrogate DNA modifications across the entire genetic loci of the previously identified genes at multiple time points – birth, 6 weeks, 12 weeks, and 36 weeks old. We identified largely sex-specific dieldrin-induced changes in DNA modifications at each time point that annotated to pathways important for neurodevelopment, potentially related to critical steps in early neurodevelopment, dopaminergic neuron differentiation, synaptogenesis, synaptic plasticity, and glial-neuron interactions. Despite large numbers of age-specific DNA modifications, longitudinal analysis identified a small number of DMCs with dieldrin-induced deflection of epigenetic aging. The sex-specificity of these results adds to evidence that sex-specific responses to PD-related exposures may underly sex-specific differences in disease. Overall, these data support the idea that developmental dieldrin exposure leads to changes in epigenetic patterns that persist after the exposure period and disrupt critical neurodevelopmental pathways, thereby impacting risk of late life diseases, including PD.

## Introduction

Parkinson’s disease (PD) is the most common neurodegenerative movement disorder and one of the fastest growing neurological diseases worldwide(Yang, W. *et al*., 2020,Dorsey, E. Ray *et al*., 2018,Dorsey, E. R. *et al*., 2007,Dorsey, E. Ray, Sherer *et al*., 2018,Willis, A. W. *et al*., 2022,Marras *et al*., 2018). PD diagnoses in the US cluster in regions with a history of industrialization – e.g. the Midwest and Northeast – and worldwide rates of disease are increasing most rapidly in newly industrialized regions, suggesting that environmental factors related to industrialization play an important role in PD etiology (Willis, A. Wright *et al*., 2010,Willis, A. W. *et al*., 2022,Dorsey, E. R. *et al*., 2007). In addition, multiple animal studies have shown that exposure to specific environmental toxicants, including certain industrial toxicants, heavy metals, and pesticides, is associated with increased risk of PD (Freire and Koifman, 2012,Caudle *et al*., 2012,Goldman *et al*., 2017,Goldman, 2014,Moretto and Colosio, 2011,Cicchetti *et al*., 2009).

Further supporting the idea that environment plays a role in PD etiology, the vast majority (90-95%) of PD cases are sporadic, with monogenic mutations responsible for only 5-10% of PD cases (Lill, 2016,Trinh and Farrer, 2013). In addition, heritability estimates suggest that only about a third of phenotypic variance of sporadic PD can be explained by genetics (Keller *et al*., 2012,Fernández-Santiago and Sharma, 2022). Thus, the etiology of sporadic PD is thought to involve complex interactions between aging, genetics, and environmental risk factors (Cannon and Greenamyre, 2013,Bogers *et al*., 2023,Fleming, 2017). The epigenome is recognized as a potential mediator of this relationship due to its unique sensitivity to the environment, establishment during cellular differentiation, and potential to regulate gene expression throughout the lifespan (Bianco-Miotto *et al*., 2017,Faulk and Dolinoy, 2011,Allis and Jenuwein, 2016). Given these characteristics, it is hypothesized that developmental exposures induce fixed changes in the epigenome, creating a poised epigenetic state in which exposure programs a modified response to later-life challenges resulting in altered disease risk (Svoboda *et al*., 2022a). A growing body of work suggests that the epigenetic mechanisms serve as a mediator of environmental risk factors in PD (Schaffner and Kobor, 2022,Tsalenchuk *et al*., 2023,Gionco and Bernstein, 2024).

According to the developmental origins of health and disease (DoHAD) hypothesis, exposures, even during prenatal development, can produce long-lasting changes in gene regulation and neurodevelopment that contribute to the risk of later-life disease (Heindel and Vandenberg, 2015,Hochberg *et al*., 2011). In such a model, the toxicant-induced mechanisms than contribute to PD risk may be temporally separated from disease onset and represent early pre-degenerative changes. Two-hit models of PD, including the developmental dieldrin α-synuclein preformed fibril (α-syn PFF) model developed in our lab, offer an opportunity to explore effects of environmental exposures and to identify pre-degenerative changes that occur prior to onset of neurodegeneration and set the stage for increased susceptibility to disease (Richardson *et al*., 2006,Gezer *et al*., 2020,Boyd *et al*., 2023). Using two-hit models, work from our lab and others has established the developmental dieldrin exposure model as a model of increased PD susceptibility (Richardson *et al*., 2006,Gezer *et al*., 2020,Boyd *et al*., 2023,Kochmanski *et al*., 2019).

Dieldrin is an organochlorine pesticide that was phased out of commercial use due to toxicological concerns in the 1970s, but the chemical persists in the environment and lipid-rich tissues like the brain due to its high stability and lipophilicity (ATSDR, 2022,Jorgenson, 2001). Mechanistic animal studies demonstrate that adult and developmental dieldrin exposures are associated with disrupted expression of PD-related proteins, oxidative stress, and increased susceptibility to secondary toxicants that affect the dopaminergic system (Hatcher *et al*., 2007,Richardson *et al*., 2006,Gezer *et al*., 2020,Kochmanski *et al*., 2019,Boyd *et al*., 2023). Specifically, adult male C57BL/6 mice developmentally exposed to dieldrin show exacerbated neurotoxicity in adulthood (12 weeks of age) induced by synucleinopathy in the α-syn PFF model and by MPTP (Gezer *et al*., 2020,Richardson *et al*., 2006,Boyd *et al*., 2023).

Previous studies have begun to explore the biological mechanisms mediating these long-lasting effects of developmental dieldrin exposure on the dopaminergic system, but these mechanisms remain incompletely defined (Richardson *et al*., 2006,Kochmanski *et al*., 2019,Gezer *et al*., 2020,Boyd *et al*., 2023). The primary mechanism of action of dieldrin is thought to be inhibition of GABA_A_ receptors (Narahashi *et al*., 1995,Narahashi, 1996). Because GABA acts as an important trophic factor in the embryonic and postnatal brain, disruption of these pathways in development is linked to neurodevelopmental and neuropsychiatric disorders; thus, dieldrin inhibition of these GABA-related pathways may affect multiple downstream pathways critical for development (Deidda *et al*., 2014). However, the mechanisms by which such inhibition leads to persistent effects on neuronal susceptibility remains incompletely defined.

Previously, we showed that developmental dieldrin exposure was associated with significant, sex-specific differential modification of cytosines (DMCs) and regions (DMRs) in genes related to dopaminergic neuron development and PD at 12 weeks of age - the time point at which MPTP or α-syn-PFFs are administered (Kochmanski *et al*., 2019). However, growing evidence suggest that environmental exposures can deflect long-term trajectories of age-related DNA modification patterns, also known as epigenetic aging (Kochmanski *et al*., 2017,Barrere-Cain and Allard, 2020). To determine if dieldrin exposure interacts with age to alter epigenetic patterns, we assessed DNA modifications at multiple time points throughout the life course – birth, 6 weeks, 12 weeks, and 36 weeks old – across the full coding regions of previously identified candidate genes using capture hybridization sequencing (Roche SeqCapEpi). The data reported here reveals potential pre-degenerative mechanisms by which early life environmental exposures contribute to late life risk of PD.

## Materials and Methods

### Animals

Adult female and male C57BL/6 mice (RRID:MGI:2159769) were purchased from Jackson Laboratory (Bar Harbor, Maine). Seven-week-old female mice were allowed to habituate after arrival for 1 week prior to beginning the developmental dieldrin exposure. Male mice were 11 weeks old upon arrival and were also allowed 1 week to habituate prior to mating. Mice were maintained on a 12-hour:12-hour reverse light/dark cycle. Mice were housed in Thoren ventilated caging systems with automatic water and 1/8-inch Bed-O-Cobs bedding with Enviro-Dri for enrichment. Food and water were available *ad libitum*. Mice were maintained on Teklad 8940 rodent diet (Envigo). After arrival, females were separated and individually housed during dieldrin dosing, except during the mating phase. After birth, F1 pups were group-housed by sex, with no more than 5 animals housed in each cage. No singly housed F1 animals were used in this study; as animals aged, all cages included a “buddy” littermate to ensure that even animals used for the last time point were never individually housed. All procedures were conducted in accordance with the National Institutes of Health Guide for Care and Use of Laboratory Animals and approved by the Institutional Animal Care and Use Committee at Michigan State University.

### Developmental dieldrin exposure

Dieldrin exposure was carried out as previously described and is summarized in Figure 1 (Richardson *et al*., 2006,Gezer *et al*., 2020,Kochmanski *et al*., 2019,Boyd *et al*., 2023). Adult (8-week-old) C57Bl/6 female mice (n=20 per group) were treated throughout breeding, gestation, and lactation. Following 3 days of habituation to peanut butter feeding, mice were administered 0.3 mg/kg dieldrin (ChemService) dissolved in corn oil vehicle and mixed with peanut butter pellets every 3 days (Richardson *et al*., 2006,Gezer *et al*., 2020,Kochmanski *et al*., 2019,Boyd *et al*., 2023). Control mice received an equivalent amount of corn oil vehicle mixed in peanut butter. The dieldrin dose was based on previous results showing low toxicity, but clear effects on the epigenome and neuronal susceptibility to neurotoxic insults (Richardson *et al*., 2006,Gezer *et al*., 2020,Kochmanski *et al*., 2019,Boyd *et al*., 2023). Mice were exposed via oral ingestion by the dam because the most likely route of exposure to dieldrin in humans is through ingestion of contaminated foods and ingestion of the resulting contaminated breast milk (ATSDR, 2022).

**Figure 1:**
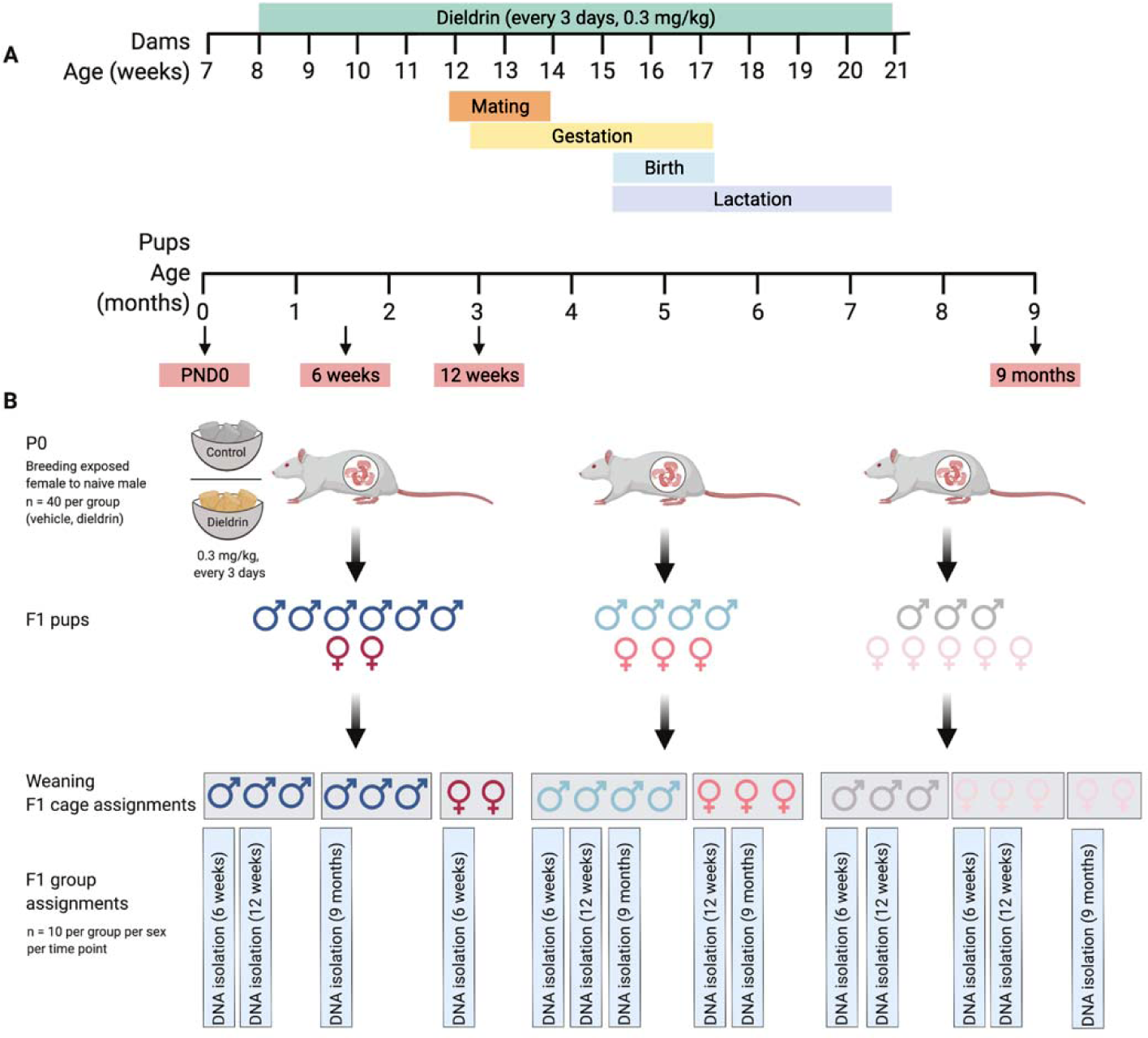
Dosing timeline, weaning strategy, and cage assignments. A) Timeline of developmental dieldrin exposure model. In this paradigm, only female F0 dams were fed dieldrin. Exposure began at 8 weeks of age with 0.3 mg/kg dieldrin dissolved in corn oil vehicle and administered via peanut butter pellets. Males were introduced for mating when females were 12 weeks of age (4 weeks into exposure). Pregnancy was confirmed by monitoring weight gain after mating. Dieldrin administration continued until pups (F1) were weaned at PND21. F1 pups were sacrificed for brain collection at birth (PND0), 6 weeks of age, 12 weeks of age, or 36 weeks of age. The birth timepoint was collected from one cohort of exposed animals, while the remaining three timepoints were collected from a second cohort of exposed animals. B) Weaning strategy and cage assignments for F1 pups followed over time. At weaning, pups (F1) were separated by sex (symbol) and litter (colors represent independent litters) with 2-5 animals per cage (grey boxes indicate cages). Animals were assigned to cages such that for each experimental group and sex, all animals were from independent litters and no animals were singly housed. In total, we generated 8-10 matched littermates per sex per group per time point. Created in BioRender.

After four weeks of exposure, unexposed C57BL/6 males (12 weeks old) were introduced for breeding for 48 hours such that male mice were not present for any peanut butter pellet feedings. To ensure adequate animal numbers for each sex at all four time points, this was carried out in two cohorts: one to generate animals for the birth time point, and one to generate animals for the 6-week, 12-week, and 36-week time points. For the birth cohort, pups were sacrificed at PND0, tissue was collected for sex determination by genotyping, and whole brains were dissected and frozen. For the remaining cohort, F1 pups were weaned and separated by litter and by sex at 3 weeks of age, with 2-5 animals per cage. At each time point, male and female littermates from independent litters were selected (*n*LJ=LJ8-10 per treatment per sex per outcome). Animals were assigned to cages such that for each experimental group and sex, all animals were from independent litters and no animals were singly housed.

The 12-week time point was selected based on previous results demonstrating increased neuronal susceptibility to MPTP and α-synuclein pre-formed fibrils (PFFs) administered at 12 weeks of age (Richardson *et al*., 2006,Gezer *et al*., 2020). The 9-month time point is equivalent to the 6-month post-PFF injection time point where we observed dieldrin-induced exacerbation of PFF-induced deficits in motor behavior and DA handling (Gezer *et al*., 2020). Birth and six weeks were selected to address the question of whether dieldrin-associated changes reflect a change in establishment or maintenance of epigenetic marks across time.

### Sex determination genotyping at birth

At the birth time point, sex determination was not possible by visual inspection and was determined using PCR for *Rmb31x/y* gametologs using an established primer pair (Tunster, 2017). DNA was isolated from tail clips taken from neonatal mice during sacrifice using the Kapa Express Extract Kit (Kapa Biosystems, Cat. #KK7100) with one minor modification – sample lysis was assisted via physical homogenization using a 1.5 mL tube plastic pestle. After lysis, samples were briefly centrifuged to pellet cellular debris, and the DNA extract was diluted 10-fold with 10 mM Tris-HCl (pH 8.0–8.5) to dilute cellular debris and digested proteins to prevent inhibition of downstream PCR. PCR was performed using 1 µL of input DNA as described (Richardson *et al*., 2006,Gezer *et al*., 2020,Kochmanski *et al*., 2019,Boyd *et al*., 2023). PCR products were visualized on a 1.2% Agarose TBE gel with ethidium bromide. During visualization, two bands of DNA were produced for male samples, and a single band of DNA was produced for female samples, allowing for accurate determination of sex at birth.

### Mass spectrometry

Dieldrin levels were measured from frozen neonatal brain tissue by the RTSF Mass Spectrometry and Metabolomics core at MSU using established methods (Hong *et al*., 2004,Anonymous 2014).

### SN microdissections

Frozen brains were mounted on a freezing cryostat (Leica, Model CM3050S) and sliced to the midbrain. Unilateral substantia nigra (SN) punches were collected using a chilled 1.0 mm micropunch and immediately placed in a frozen 1.5 mL tube on dry ice.

### DNA isolation

DNA was isolated from unilateral SN punches using Qiagen QIAamp DNA Micro Kits (Qiagen, Cat. #56304) according to the included protocols with the following minor modifications. First, prior to adding ATL buffer to SN punches, 80 µL PBS was added to each sample and a 1.5 mL tube pestle was used to break up the sample. After that, 100 µL ATL buffer was added to each homogenized sample, which were allowed to equilibrate to room temperature. Second, for the proteinase K digestion step, the 56°C incubation time was extended to overnight. Third, carrier RNA was added to Buffer AL (this is an optional step). Fourth, the incubation time after addition of 100% EtOH was increased to 10 minutes, and the incubation time for the elution step was increased to 5 minutes. Lastly, to increase yield, the elution step was repeated by reapplying elution buffer to QIAamp MinElute column. DNA was eluted in 54 µL of 10 mM Tris-HCl, pH 8.0. DNA yield and purity were determined using a Qubit 3 fluorometer (ThermoFisher) and a NanoDrop spectrophotometer (ThermoFisher). Isolated DNA was stored at −80°C prior to sequencing.

### Selection of candidate regions

To select candidate regions, we targeted all intragenic genomic features and known enhancers at genes annotated to DMCs or DMRs in our previous study (Kochmanski *et al*., 2019). Targeted regions include 255 male-specific annotations and 1043 female-specific annotations and represent a total of ∼11.4 Mb of genomic space (Supplementary File 1). We included regions identified for both male- and female-specific changes to determine if these are truly sex-specific. Intergenic regions annotated to our candidate genes account for 151 Mb of genomic space, making inclusion cost-prohibitive with the SeqCapEpi platform. In addition, these intergenic regions remain largely unexplored, making data generated from these regions of limited interpretability. SeqCapEpi custom bait probes for regions of interest were designed using the Roche NimbleDesign software (Supplementary File 2). Baits covered 91.6% of target bases for a total capture space of ∼10.4 Mb.

### Capture hybridization sequencing

Capture hybridization-sequencing libraries were prepared by the Van Andel Genomics Core from 100 ng of high molecular weight DNA using the Accel-NGS Methyl-Seq DNA Library kit (v3.0) (Swift Biosciences, Cat. #30024). DNA was sheared following manufacturer’s protocol to an average size of 250bp, and sheared DNA was bisulfite converted using the EZ DNA Methylation-Gold kit (Zymo Research, Cat. #D5005) with an elution volume of 15 µL. Following adapter ligation, 8 cycles of library amplification were performed. Amplified libraries were pooled in batches of 8 and targeted enrichment of a custom 11Mb region was performed using Roche SeqCapEpi developer probes and SeqCapHyperEpi workflow starting at step 4.0 with the following modification: the capture was performed using IDT xGen Universal blockers to replace the SeqCap HE Universal Oligo and SeqCap HE Index Oligo. The post-capture amplification was also adjusted to 11 cycles of amplification and the final extension changed from 30 seconds to 1 minute. Quality and quantity of the finished library pools were assessed using a combination of Agilent DNA High Sensitivity chip (Agilent Technologies, Inc.) and QuantiFluor® dsDNA System (Promega Corp.). Sequencing (100 bp, paired end) was performed on an Illumina NovaSeq6000 sequencer using an S4, 200 bp sequencing kit (Illumina Inc.) with 10% PhiX included to improve base diversity. Base calling was done by Illumina Real Time Analysis 3 and output of NextSeq Control Software was demultiplexed and converted to FastQ format with the Illumina *Bcl2fastq* software (version 1.9.0).

### Data processing

FastQ files were processed using a slightly modified form of the bioinformatics pipeline previously established by our group to analyze reduced representation bisulfite sequencing (RRBS) data (Richardson *et al*., 2006,Gezer *et al*., 2020,Kochmanski *et al*., 2019,Boyd *et al*., 2023). Command line tools and the open-source statistical software R (version 4.1.2) were used for all analyses. For all sequencing data, the *FastQC* tool (version 0.11.7) was used for data quality control, and the *trim_galore* tool (version 0.4.5) was used for adapter trimming (Andrews, 2016,Krueger, 2017). During adapter trimming, we used the default minimum quality score and added a stringency value of 6, thereby requiring a minimum overlap of 6 bp. Trimmed CapHyb-seq reads were aligned to the *mm10* reference genome using *bismark* (version 0.19.1) (Krueger and Andrews, 2011). Methylation data were extracted from the aligned reads in *bismark* using a minimum threshold of 5 reads to include a CpG site in analysis.

### Verification of sequencing coverage

To determine the sequencing coverage across the targeted regions, we used BedTools Basic Protocol 3 for measuring coverage in targeted DNA sequencing experiments as described (Quinlan, 2014). BAM files for each sample obtained from *bismark* were compared with targeted bases (Supplementary File 2) using the *coverage* and *multicov* functions from *bedtools* (version 2.31.0) to determine the fraction of target bases that were captured and their sequencing depths (Quinlan and Hall, 2010). In differential modification analysis, only sites with coverage ≥ 5 in all samples were included in analysis. Of 6604 target regions, 1359 remained (21%), for ∼2.8 Mb of captured sequence.

### Differential modification analysis

The *DSS* (version 2.48.0) and *DMRcate* (version 2.14.1) R packages were used to test CapHyb-seq data for differential methylation (Feng *et al*., 2014,Peters *et al*., 2015). Given that dieldrin exposure has shown sex-specific effects on the dopaminergic system, all differential modification models were stratified by sex (Kochmanski *et al*., 2019,Boyd *et al*., 2023,Richardson *et al*., 2006). All pups included in modeling were from independent litters. Of note, we refer to DNA modifications, rather than DNA methylation, since our BS-based method does not differentiate between DNA methylation and hydroxymethylation. While DNA hydroxymethylation plays a critical role in gene expression in the brain and is particularly sensitive to environmental factors, methods for differentiating these marks remain limited and cost-prohibitive, especially across multiple time points (Kochmanski and Bernstein, 2020).

### Cross-sectional differential DNA modification analysis

To test for differentially methylated cytosines (DMCs) by dieldrin exposure at each cross-sectional timepoint, we used the *DMLtest* function in *DSS* to perform two-group Wald tests. For DMLtest modeling, the equal dispersion parameter was set to *FALSE* and smoothing was set to *TRUE*. DMCs were considered significant at false discovery rate (FDR) <0.05. To test for differentially methylated regions (DMRs) at each timepoint, we combined outputs from the *callDMR* function in DSS and the *dmrcate* function in DMRcate. For *callDMR* modeling, the p-value threshold was set to 0.05, minimum length was set to 50 base pairs, and minimum CpGs was set to 3. Meanwhile, for *dmrcate* modeling, the lambda value was set to 500, the C value was set to 4, and minimum CpGs was set to 3. DMR significance for the *dmrcate* output was set to a minimum smoothed FDR < 0.05.

### Longitudinal differential DNA modification analysis

To test for DMCs by age, as well as simultaneous age and exposure in a multivariate model, we used the *DMLfit.multiFactor* function in *DSS* to perform linear models using a general experimental design. Given that age was an ordered variable, we coded each age group as a number for modeling – 6 weeks old = 1, 12 weeks old = 2, and 36 weeks old = 3. The birth timepoint was not included in longitudinal models because the F1 offspring came from a separate cohort of exposed animals, meaning they were not matched littermates like the later three time points. In the model for age only, coded age was included as the only independent variable. Meanwhile, in the multivariate model, exposure was included as a two-group categorical variable (“dieldrin”, “control”), and age:exposure was included as an interaction term. For the age alone and age:exposure models, DMCs were considered significant at false discovery rate (FDR) <0.05.

### Genomic annotation

After differential methylation testing, the *annotatr* R package (version 1.26.0) was used to annotate identified DMCs and DMRs to the reference *mm10* genome (Cavalcante and Sartor, 2017). Within *annotatr*, the *annotate_regions* function was used to generate CpG context, gene body, and regulatory feature annotations. Annotations for miRNA (miRbase), ENCODE predicted mouse midbrain enhancers (Accession: ENCSR114ZIJ), and custom full stack ChromHMM chromatin states for *mm10* databases were added to the annotation cache in *annotatr* (Vu and Ernst, 2023,Kozomara *et al*., 2019,Luo *et al*., 2020,Kundaje *et al*., 2012).

### Data visualization

Raw CapHyb-seq beta values were extracted using the *bsseq* R package (1.36.0). Volcano plots were generated using the *ggplot2* R package (version 3.4.4). Venn and Euler diagrams were generated with the *eulerr* R package (version 7.0.1) (Larsson, 2024). UpSet plots were generated using the *UpSetR* package (version 1.4.0) (Conway *et al*., 2017). The *ComplexHeatmap* R package (version 2.18.0) was used to visualize beta value differences over time for all DMCs significant at one or more of the four time points. Specific genes of interest were visualized using the WASHU Epigenome Browser to determine genomic location and context of DM regions; chromHMM annotations were loaded to compare DM loci with chromatin state annotations (Conway *et al*., 2017,Li *et al*., 2022). The R packages *ggplot2*, *ggpubr* (version 0.6.0), and *ggeasy* (version 0.1.4) were used to generate violin plots to display raw beta values for premature aging DMCs (Carroll *et al*., 2023,Kassambara, 2023).

### Pathway and Network Analysis

Gene ontology (GO) term enrichment testing and pathway analysis was performed on genes annotated to male and female DMCs and DMRs using the ClueGO application in Cytoscape (version 3.10.1) (Bindea *et al*., 2009,Smoot *et al*., 2011).

For *ClueGO* testing, genes annotated to DMCs and DMRs stratified by sex and timepoint were input as separate gene lists, “groups” was selected as the visual style, and the GO-biological process (GOBP) term was included for enrichment testing. Network specificity was set to “Medium”, such that the GO Tree Interval minimum was equal to 3 and the maximum was equal to 8. Only terms with at least 3 genes and a Bonferroni-corrected p-value < 0.05 were included in pathway visualizations. The connectivity score (Kappa) was set at 0.4, and default GO Term Grouping settings were used in all analyses. The genes found in enriched GOBP terms by *ClueGO* were used for STRING network analysis and to create Venn diagrams for each sex displaying the overlap at different time points.

Protein-protein interaction network analysis was performed using STRING (version 12.0) with the genes found in enriched GOBP terms by ClueGO as input (Szklarczyk *et al*., 2017,Szklarczyk *et al*., 2015). STRING network analysis was performed using default parameters, including a minimum required interaction score = 0.4 and all interaction sources activated.

All code and metadata used for these analyses is provided in Supplementary Files 13-20.

## Results

### Cross-sectional analysis of DNA modifications

In our first stage of analysis, we stratified data by sex and identified significant DMCs and DMRs at each assessed time point. DMCs and DMRs results are summarized in Table 1 and DMCs are summarized in Figure 2a-c. The large majority of the intragenic DM loci are found within introns, as expected based on genomic space (Figure 2e). The majority of DM loci are found in regions annotated by chromatin state analysis as active enhancers, with weak enhancers and promoters as the next two most frequent categories (Figure 2f). Complete cross-sectional results and annotations are included in Supplementary Files 3-7. The numbers of genes containing DMCs or DMRs were determined after annotation and are summarized in Table 2. There are far fewer genes containing DMCs and DMRs indicating that each gene contains multiple sites of differential modification (Figure 2c,d). To generate gene lists for subsequent analysis, we identified unique gene lists for each sex and time point (Table 2, Supplementary File 8).

**Figure 2:**
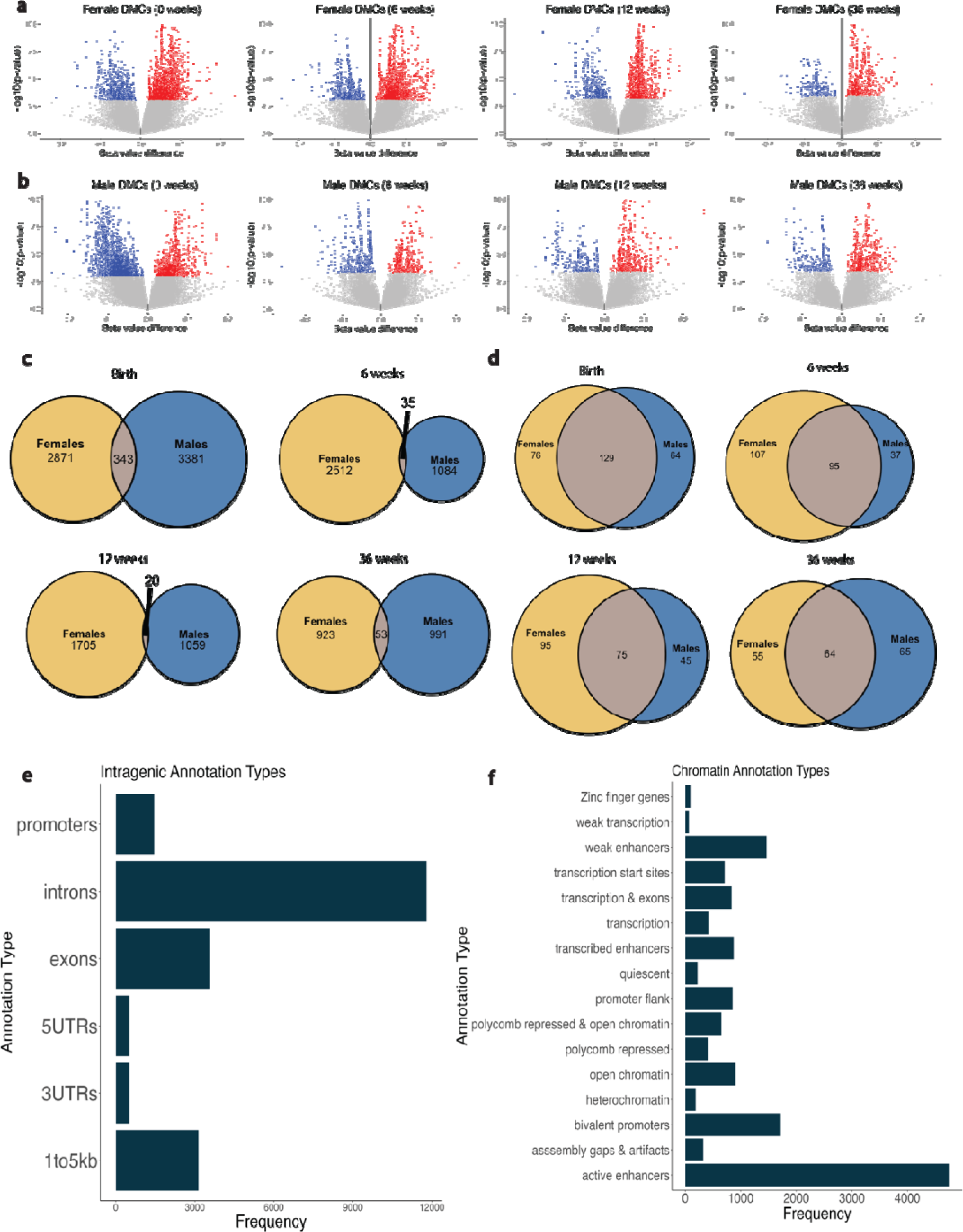
Significant DMCs for each sex at each time point. a,b) Volcano plots of DMCs for (a) female animal,and (b) male animals at each time point. Beta value differences, plotted on the x-axis, represent the difference between mean methylation values for the control groups from the mean methylation values for the dieldrin groups, such that positive values indicate hypermodified cytosines in exposed animals and negative values indicate hypomodified cytosines in exposed animals. Colored/dark points represent statistically significant DMCs (FDR<0.05). The blue points represent hypomodified DMCs and the red points represent hypermodified DMCs in dieldrin animals versus control animals. c,d) Euler diagrams display overlap between sexes at each time of (c) significant DMCs by location or (d) genes containing DMCs or DMRs. e,f) Frequency histograms of DMC and DMR annotations for male and female data combined for (e) intragenic annotation types (f) and chromatin annotation types. Annotation numbers are higher the DMC/DMR numbers because most DMC/DMRs have multiple annotations.

**Table 1:** Numbers of identified DMCs and DMRs in each sex at each time point.

|  | DMCs |  | DMRs |  |
| --- | --- | --- | --- | --- |
|  | Female | Male | Female | Male |
| **Birth** | **3214** | **3724** | **152** | **248** |
| Hypermodification | 2335 | 1212 | 112 | 59 |
| Hypomodification | 879 | 2512 | 40 | 189 |
| **6 weeks** | **2547** | **1119** | **130** | **53** |
| Hypermodification | 1834 | 577 | 97 | 23 |
| Hypomodification | 713 | 542 | 33 | 30 |
| **12 weeks** | **1725** | **1079** | **73** | **44** |
| Hypermodification | 1177 | 688 | 42 | 26 |
| Hypomodification | 548 | 391 | 31 | 18 |
| **36 weeks** | **976** | **1044** | **31** | **46** |
| Hypermodification | 703 | 718 | 19 | 28 |
| Hypomodification | 273 | 326 | 12 | 18 |

**Table 2:** Numbers of unique genes annotated to DMRs and DMCs. DMC Genes and DMR Genes columns include unique genes in each category. Many genes have multiple sites of differential modification. The Unique Genes column indicates the number of unique genes annotated to DMCs and DMRs for each sex and time point. These final lists of unique genes for each time point were utilized in downstream analysis steps.

|  | DMC Genes |  | DMR Genes |  | Unique Genes |  |
| --- | --- | --- | --- | --- | --- | --- |
|  | Female | Male | Female | Male | Female | Male |
| **Birth** | **204** | **188** | **91** | **95** | **205** | **193** |
| Hypermodification | 154 | 134 | 63 | 46 |  |  |
| Hypomodification | 111 | 133 | 35 | 72 |  |  |
| **6 weeks** | **200** | **128** | **84** | **43** | **202** | **132** |
| Hypermodification | 148 | 78 | 61 | 20 |  |  |
| Hypomodification | 105 | 82 | 29 | 26 |  |  |
| **12 weeks** | **166** | **116** | **64** | **38** | **170** | **120** |
| Hypermodification | 116 | 93 | 42 | 26 |  |  |
| Hypomodification | 94 | 52 | 29 | 16 |  |  |
| **36 weeks** | **119** | **126** | **32** | **36** | **119** | **129** |
| Hypermodification | 85 | 90 | 21 | 24 |  |  |
| Hypomodification | 62 | 68 | 11 | 17 |  |  |

We also identified genes that containing DMCs or DMRs at all time points for each sex (Figure 4). For all genes, 63 and 45 genes were differentially modified at all time points in female and male animals, respectively, with 32 shared genes between sexes at all time points (Figure 4a,b). For genes included in enriched GO terms, only 8 genes were differentially modified at all time points for female animals and only 8 for male animals. Only 6 were differentially modified at all time points in both sexes (*Fgfr2, Myo3b, Ephb2, Ptk7, Prkca, Ppp1r16b*) (Figure 4c,d). Genes differentially modified at all time points in male animals, but not female animals, include *Gnas* and *Prkce*, which are differentially modified in female animals at 6, 12 and 36 weeks, and 6 and 12 weeks, respectively. Genes differentially modified at all time points in female animals, but not male animals include *Rhoq* and *Xylt1*, which are differentially modified in male animals at 6 weeks, and birth and 6 weeks, respectively. At each time point for each of these genes, the location of DMCs and DMRs are largely inconsistent, indicating that the differential modification of these genes is complex and dynamic with sex, age, and location-specific changes.

**Figure 3:**
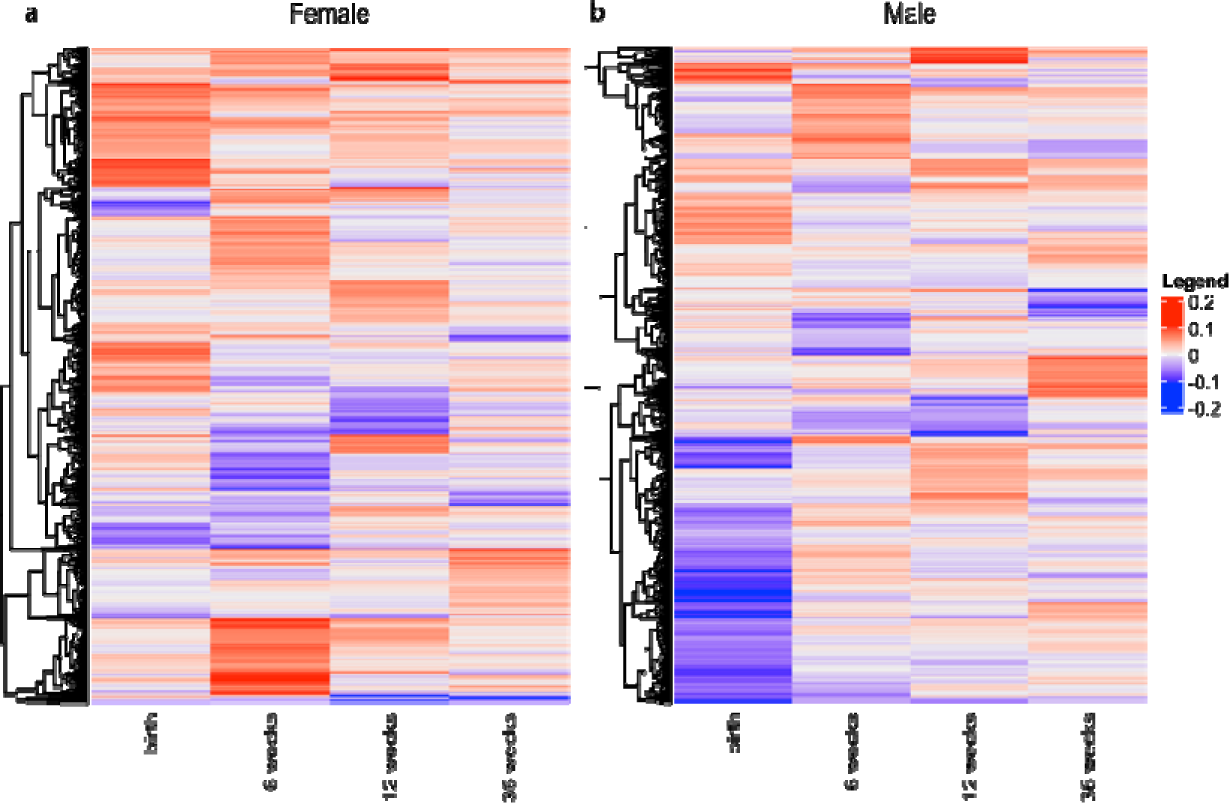
Beta value differences across cross-sectional time points. Heatmaps display the beta value differences for significant DMCs over time for (a) female animals and (b) male animals. Each row represents a single DMC. Blue indicates hypomodification in dieldrin exposed animals compared to control and red indicates hypermodification in dieldrin exposed animals compared to control. Rows are clustered based on similarity of beta value differences, with DMCs hypomodified at all time points at the bottom and those hypermodified at all time points at the top.

**Figure 4:**
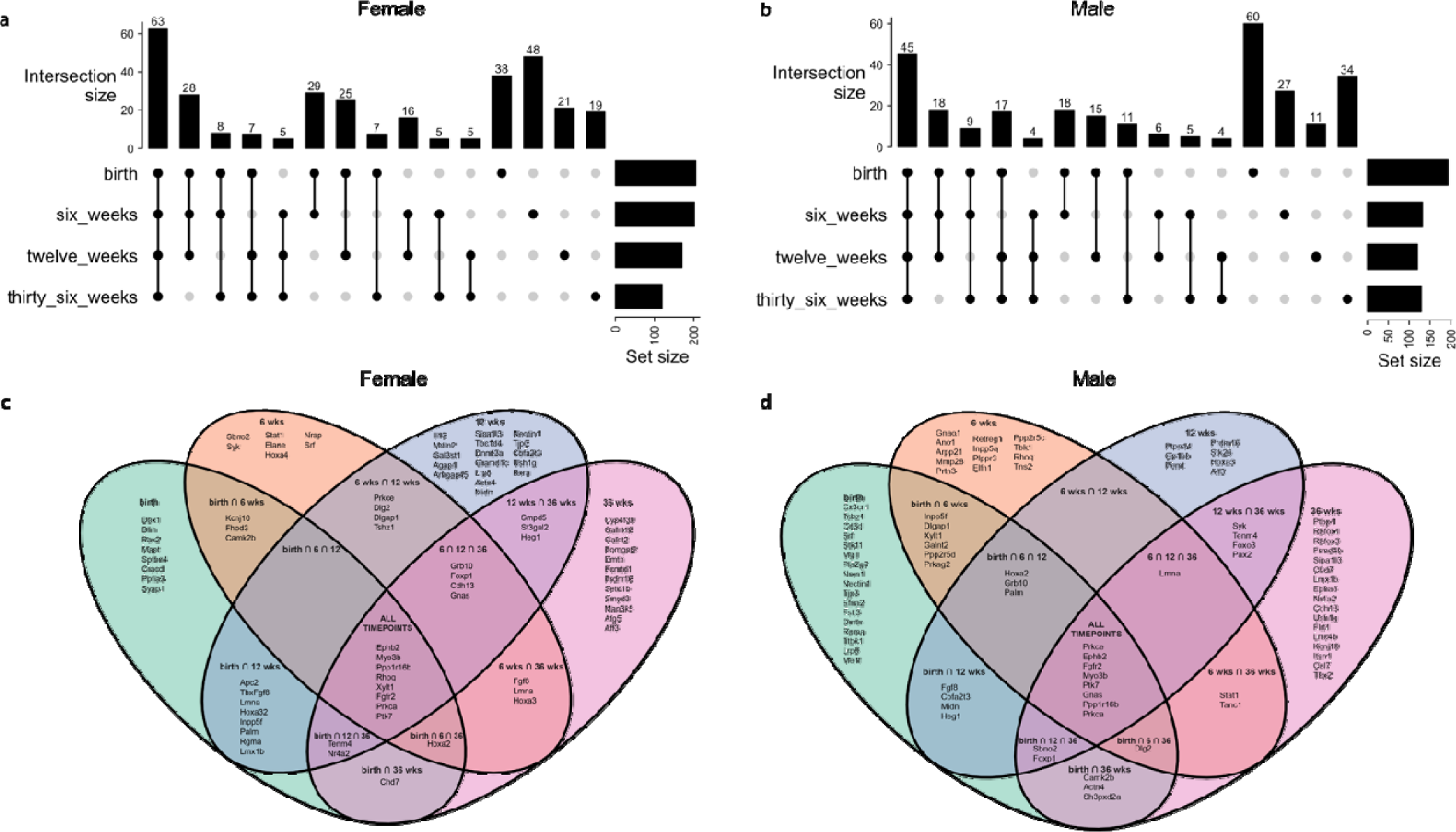
Overlap between differentially modified genes in female and male animals at cross-sectional time points. (a,b) UpSet plots show overlap between all differentially modified genes at each time point. (c,d) Venn diagrams visualize the overlap between the genes found in enriched GO terms at each time point.

To explore if there are clear patterns over time by specific location, we generated heatmaps of all Significant DMCs clustered by patterns in the direction of change at each time point, with DMCs hypomodified at all time points at the bottom and those hypermodified at all time points at the top (Figure 3). We observed an overall decline in the number of both DMCs and DMRs with increasing age in both male and female animals (Figure 2a,b, Table 1). In addition, we observed a greater number of DMCs/DMRs with increased modification at birth in female animals, but a greater number of DMCs/DMRs with decreased modification at birth in male animals. At the DMC level, this pattern remains for female animals at later time points but reverses for male animals by 6 weeks for DMCs and by 12 weeks for DMRs. Heatmaps show changes in beta value difference over time, but there is not an overall, consistent indication at this level of analysis that dieldrin is causing global deficits in establishment or maintenance of epigenetic marks or an acceleration epigenetic aging (Figure 3).

### Functional annotation of differentially modified genes

To determine if differentially modified (DM) genes function in known pathways and networks, we generated unique gene lists from annotated DMC and DMR data for each sex and time point for downstream analysis (Supplementary File 8, Table 2). From each list, gene ontology term enrichment analysis was performed in ClueGO (Supplementary Tables 1 and 2). Genes within enriched GO terms were used as input for STRING network analysis to identify potential functional interactions between genes. Because the genes in these networks were overlapping across timepoints, networks for all time points group together are shown in Figure 5, while networks for each separate time point are shown in Supplementary Figure 1. For female data, of the 79 genes in enriched GO terms, 42 genes were included in a highly interconnected network. Similarly, for male data, of the 88 genes in enriched GO terms, 50 genes were included in a highly interconnected network. That these genes are highly interconnected was expected since these genes were selected for this analysis based on shared GO terms and potential interactions in STRING in our previous study (Kochmanski *et al*., 2019).

**Figure 5:**
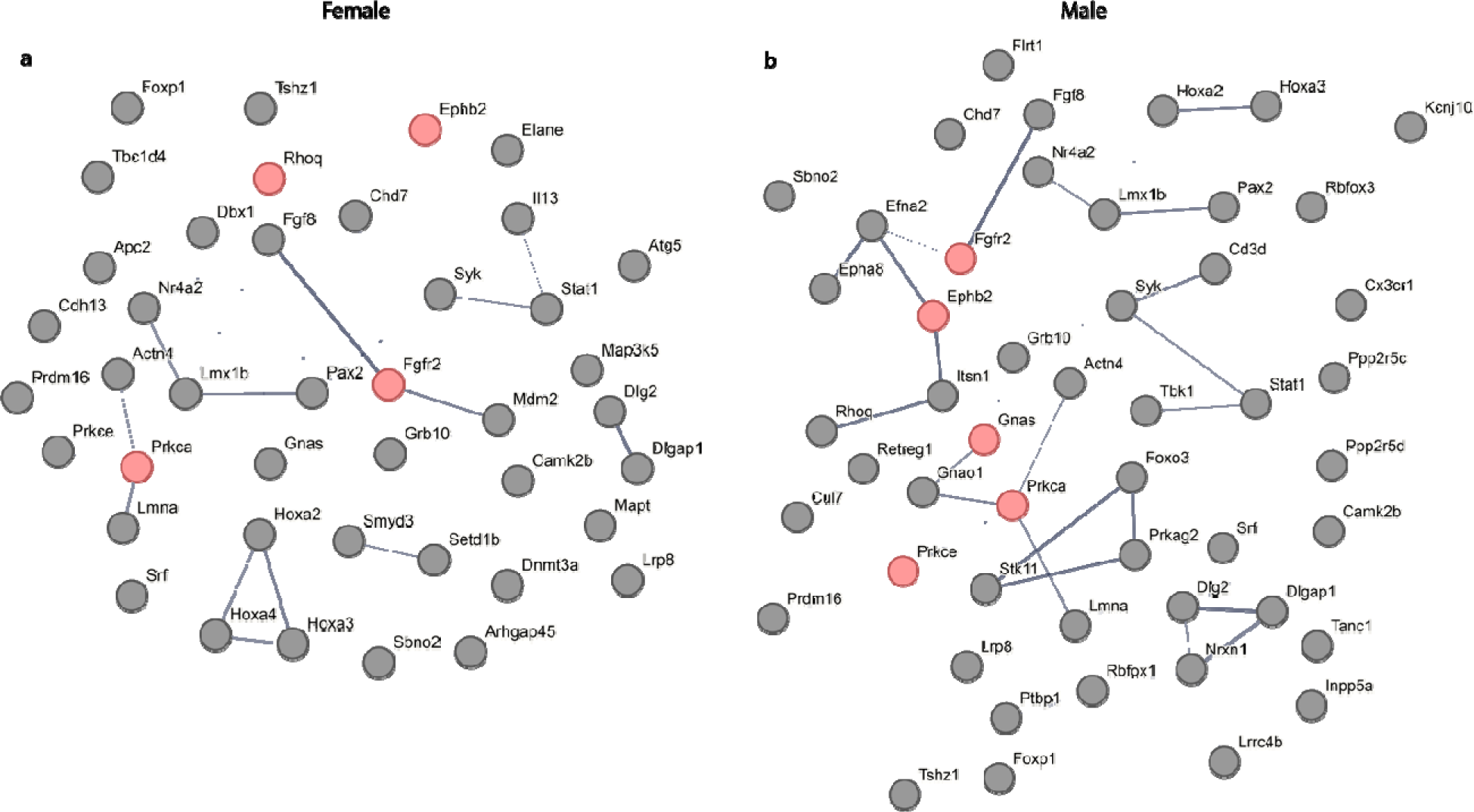
STRING protein-protein interaction networks between genes found in enriched GO terms. Interaction networks for DM genes from female (a) and male (b) animals at all time points combined. STRING networks display the interactions with at least a 0.4 interaction score and omit any disconnected nodes. Each node represents all proteins produced by a single, protein coding gene locus. Edges represent protein-protein associations, but not necessarily physical interactions. Thickness of the lines indicates the strength of data support. Genes identified at all time points are highlighted in red.

### Selection and characterization of candidate genes

From the large list of DM genes, we selected candidate genes to highlight with related functions in key pathways based on confirmed expression in midbrain/SN, connections in GO term and STRING analyses, and *a priori* knowledge of these genes (Figure 5, Table 3, Supplementary Tables 1,2). These include genes involved in dopamine neurogenesis and the differentiation and survival of midbrain DA neurons (*Nr4a2, Lmx1b*); fibroblast growth factor signaling (*Fgf8, Fgfr2, Stat1*); and synaptogenesis, maintenance of synaptic structure, and synaptic plasticity (*Ephb2*, *Nrxn1*, *Dlg2*, *Dlgap1*, *Camk2b*, *Prkca*, *Prkce*) and two imprinted genes (*Gnas, Grb10)*.

**Table 3:** Summary of selected differentially modified genes. Genes containing DMCs and DMRs that are described in text are listed, with asterisks indicating the time points at which DMCs or DMRs were identified. Genes visualized in Figure 6 are indicated in bold. Cell-type expression in the substantia nigra from the Allen Brain Cell Atlas are indicated, with +++ indicating high relative expression compared to other cell types, ++ indicating lower relative expression compared to other cell types, and + indicating sparse expression in only a few cells. Enriched GO terms indicate those that contained the gene in our ClueGO enrichment analysis. Abbreviations: 0 = PND0, 6 = 6 weeks, 12 = 12 weeks, 36 = 9 months, Neu = dopamine neurons, MG = microglia, Astro = astrocytes, oligo = oligodendrocytes.

| Gene | Female |  |  |  | Male |  |  |  | SN expression |  |  |  | Enriched GO Terms |
| --- | --- | --- | --- | --- | --- | --- | --- | --- | --- | --- | --- | --- | --- |
|  | 0 | 6 | 12 | 36 | 0 | 6 | 12 | 36 | Neu | MG | Astro | Oligo |  |
| Dopamine neurogenesis, differentiation and survival of midbrain DA neurons |  |  |  |  |  |  |  |  |  |  |  |  |  |
| Lmx1b | * |  | * |  |  |  |  | * | +++ | + | + | + | central nervous system neuron development, dopaminergic neuron differentiation |
| Nr4a2 | * |  | * | * |  |  |  | * | +++ | + | + | + | central nervous system neuron development, dopaminergic neuron differentiation, adult locomotory behavior, central nervous system neuron axonogenesis |
| Fibroblast growth factor signaling |  |  |  |  |  |  |  |  |  |  |  |  |  |
| Fgf8 |  | * |  | * | * |  | * |  | + | + | + | + | ear morphogenesis, lung morphogenesis, motor neuron axon guidance, cranial skeletal system development, animal organ formation, specification of animal organ identity, positive regulation of stem cell proliferation, hindlimb morphogenesis, inner ear morphogenesis, brain morphogenesis, embryonic skeletal system morphogenesis, embryonic cranial skeleton morphogenesis, embryonic hindlimb morphogenesis |
| Fgfr2 | * | * | * | * | * | * | * | * | +++ | +++ | +++ | +++ | ear morphogenesis, lung morphogenesis, mesenchymal cell proliferation, regulation of mesenchymal cell proliferation, positive regulation of mesenchymal cell proliferation, cellular response to retinoic acid, embryonic cranial skeleton morphogenesis, cardiac muscle cell proliferation, regulation of cardiac muscle cell proliferation, cranial skeletal system development, animal organ formation, specification of animal organ identity, positive regulation of stem cell proliferation, inner ear morphogenesis, embryonic skeletal system morphogenesis, embryonic cranial skeleton morphogenesis, striated muscle cell proliferation, positive regulation of smooth muscle cell proliferation, central nervous system neuron development, lipopolysaccharide-mediated signaling pathway |
| Stat1 | * | * |  |  | * | * |  | * | +++ | +++ | +++ | +++ | regulation of mesenchymal cell proliferation, positive regulation of smooth muscle cell proliferation, lipopolysaccharide-mediated signaling pathway, mesenchymal cell proliferation, tumor necrosis factor-mediated signaling pathway, syncytium formation, syncytium formation by plasma membrane fusion, positive regulation of mesenchymal cell proliferation |
| Synaptogenesis, maintenance of synaptic structure, neurotransmission, and synaptic plasticity |  |  |  |  |  |  |  |  |  |  |  |  |  |
| Ephb2 | * | * | * | * | * | * | * | * | +++ | + | + | +++ | ear morphogenesis, regulation of neurotransmitter receptor activity, filopodium assembly, regulation of filopodium assembly, regulation of cell morphogenesis involved in differentiation, cranial nerve morphogenesis, cranial nerve development, inner ear morphogenesis, positive regulation of synapse assembly, central nervous system neuron development, central nervous system neuron axonogenesis, regulation of neuronal synaptic plasticity, regulation of long-term neuronal synaptic plasticity, positive regulation of dendritic spine development, positive regulation of dendrite morphogenesis |
| Dlg2 | * | * | * | * | * | * |  | * | +++ | + | +++ | +++ | structural constituent of postsynapse, protein localization to cell junction |
| Dlgap1 |  | * | * |  | * | * | * |  | +++ | + | +++ | +++ | structural constituent of postsynapse, protein localization to cell junction |
| Nrxn1 |  |  |  |  | * |  |  |  | +++ | + | +++ | +++ | synapse maturation, regulation of neurotransmitter receptor activity, protein localization to cell junction, filopodium assembly, regulation of filopodium assembly |
| Camk2b | * | * |  |  | * |  |  | * | +++ | + | +++ | +++ | regulation of neuronal synaptic plasticity, structural constituent of postsynapse, regulation of neuronal synaptic plasticity |
| Prkca | * | * | * | * | * | * | * | * | +++ | +++ | +++ | +++ | central nervous system neuron development, central nervous system neuron axonogenesis, negative regulation of glucose import, positive regulation of smooth muscle cell proliferation, positive regulation of smooth muscle cell proliferation, central nervous system neuron development, negative regulation of glucose transmembrane transport, platelet aggregation |
| Prkce | * | * | * | * | * | * | * | * | +++ | +++ | +++ | +++ | macrophage activation involved in immune response, lipopolysaccharide-mediated signaling pathway, chemosensory behavior |
| Imprinted Genes |  |  |  |  |  |  |  |  |  |  |  |  |  |
| Grb10 | * | * | * | * | * | * | * | * | +++ | + | + | + | negative regulation of carbohydrate metabolic process, negative regulation of cellular response to insulin stimulus, negative regulation of glucose transmembrane transport, negative regulation of insulin receptor signaling pathway, regulation of glucose import, negative regulation of glucose import, negative regulation of cellular carbohydrate metabolic process, positive regulation of cold-induced thermogenesis |
| Gnas | * | * | * | * | * | * | * | * | +++ | +++ | +++ | +++ | embryonic skeletal system development, embryonic skeletal system morphogenesis, cellular response to alcohol, platelet aggregation, cranial skeletal system development, hindlimb morphogenesis, embryonic cranial skeleton morphogenesis, embryonic hindlimb morphogenesis |
| Epigenetic regulators |  |  |  |  |  |  |  |  |  |  |  |  |  |
| Dnmt3a | * | * | * |  | * |  |  |  | +++ | +++ | +++ | +++ | cellular response to alcohol |
| Hdac9 | * | * | * | * | * | * | * | * | +++ | +++ | +++ | +++ | Not found in enriched GO terms |

Overall, we found that candidate gene DMCs and DMRs tend to cluster together and that hypomodified and hypermodified DMCs and DMRs occur together (i.e. these are not interspersed within clusters of differential modifications). Highlighting the sex-specificity and age-dependence of epigenetic regulation, there was limited overlap within these genes between timepoints and sexes in location or direction of change, even in those genes that were identified in both sexes and all timepoints. In addition, consistent with the frequency of DMC/DMR annotations, identified DM loci occur in regions likely to be regulated by differential modifications, including regions annotated by chromHMM as promoters, transcription start sites of major and alternate transcripts, and enhancers (Figure 2,Figure 6, Supplementary Files 6,7).

**Figure 6:**
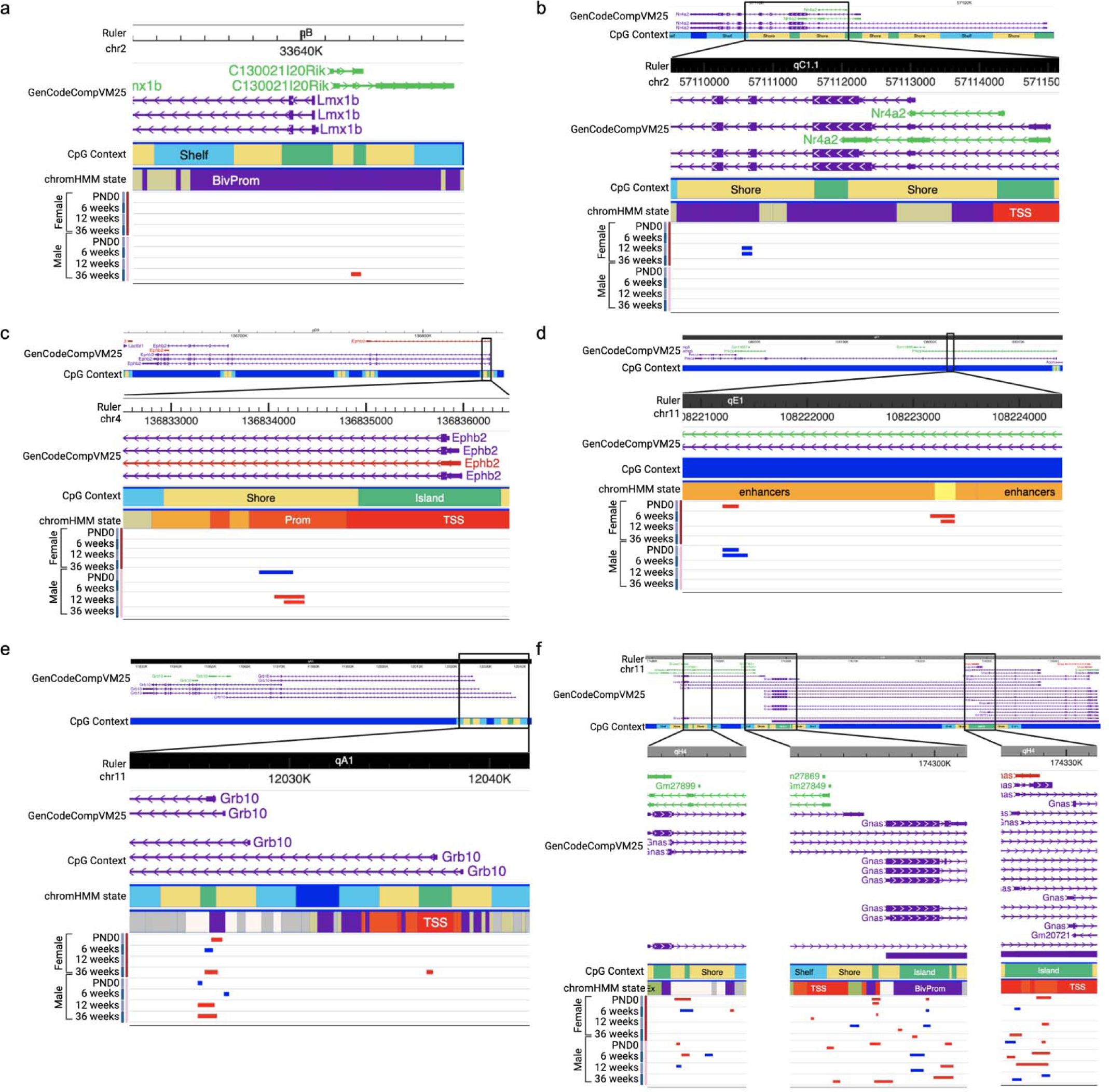
Visualization of target genes in the Wash U Epigenome Browser. Selected DMRs were visualized in the Wash U Epigenome browser showing CpG context and a ChromHMM track generated from the universal chromatin state annotation for *mm10* (Vu and Ernst, 2023). For b-f, the full genomic locus is shown above; the box indicates the location of the zoomed in region below. (a) A male-specific hypermodified DMR annotated to *Lmx1b* in a bivalent promoter (purple). (b) A female-specific hypomodified DMR annotated to *Nr4a2* spans an exon-intron boundary and maps to a region annotated as a bivalent promoter (purple). (c) Male-specific overlapping DMRs in *Ephb2* are found in a promoter flanking region (orange); the direction of change is different at birth and 12 weeks. (d) A DMR within intron 2 of *Prkca* maps to and active enhancer (light orange) and the direction of change is opposite in male and female samples. A neighboring female-specific hypermodified DMR was identified at 6 weeks only. (e) DM loci annotated to *Grb10* are located within two neighboring CpG islands in a bivalent promoter (purple) and transcription starts site (TSS; red) in a highly sex-, age- and location specific manner. (f) DM regions in *Gnas* are located in all 3 CpG islands/promoters of this gene with no consistent pattern in direction of change by age or location. ***Color coding for GenCode annotation:*** Purple – coding, Green – non-coding, Red – problem transcript. ***Color coding for CpG context:*** Green – CpG island, yellow – shore, light blue – shelf. ***Color coding for chromHMM state:*** Purple – bivalent promoter; bright orange – promoter flanking, light orange – active enhancer, red – transcription start site. ***Color coding for DMRs:*** Blue indicates hypomodification; red indicates hypermodification.

In Figure 6, we used the WashU Epigenome Browser to explore genomic location and context of DMCs and DMRs for the candidate genes. Selected regions are described below and were visualized in the WashU Epigenome Brower to highlight examples of the sex-, age-, and location-specificity of dieldrin-induced differential modifications (Figure 6)(Li *et al*., 2022).

### Sex-specific differential modifications

A male-specific hypermodified DMR found only at 9 months annotated to *Lmx1b* is located in a bivalent promoter that drives expression of *Lmx1b* and C130021l20Rik, a co-expressed lncRNA (Figure 6a). Multiple DM locie annotated to *Nr4a2* was identified in female animals at PND0, 12 weeks and 9 months and in male animals at 9 months only. Shown is a female-specific hypomodified DMR in *Nr4a2* found at 12 weeks only that spans an exon-intron boundary and maps to a region annotated as a bivalent promoter (Figure 6b). DM loci annotated to *Prkca* within were identified in both sexes at all time points. These map to intron 2 and overlap with a non-coding RNA within this locus. The DMR shown maps to an active enhancer, was found at birth in both sexes, and is hypermodified in female but hypomodified in in male samples (Figure 6d).

### Age-specific differential modifications

DMC/DMRs were identified at all timepoints in both sexes in *Ephb2,* and they all map to intron 1. Male-specific overlapping DMRs found in a promoter flanking region including the one male hypermodified DMR overlap by location, but the direction of change is different at birth and 12 weeks of age (Figure 6c).

### Complex differential modification of imprinted genes

Differential modification of these imprinted genes are highly age-, sex- and location-specific.DM loci annotated to *Grb10* were identified in both sexes at all time points and all are located within two neighboring CpG islands. One hypermodified DMR in female samples at 36 weeks is located within the transcription start site. The downstream CpG island contains multiple overlapping DM regions within a bivalent promoter at birth (hyper), 6 weeks (hypo), and 9 months (hyper) in female samples, and at all and 9 months). DM regions were identified in the imprinted gene *Gnas* at all time points in both sexes. These are located in all 3 CpG islands/promoters of these gene with no consistent pattern in direction of change by age, sex, or location.

In Table 3, we summarize data on these selected genes, including the time points and sex at which each contained DM loci, cell type expression within the SN from the Allen Brain Cell Atlas, and the GO terms each gene mapped to in the ClueGO analysis (Yao *et al*., 2023). Because the current data is derived from a bulk tissue micropunch, we do not have cell-type specific data. Thus, we cross-referenced candidate genes to the Allen Brain Cell Atlas to identify genes with known SN expression in mice 7-10 weeks old, as well as which cell types they are known to be expressed in (Yao *et al*., 2023).

### Dieldrin-induced deflection of age-related DNA modification patterns

To determine whether dieldrin deflects long-term trajectories of age-related DNA modification patterns, we first identified DMCs with age-related changes in only control animals at the three time points where data were collected from matched littermates across time – 6 weeks, 12 weeks, and 36 weeks old. We identified 290 age-related DMCs in males and 408 age-related DMCs in females (Table 4). Consistent with the cross-sectional analyses, these DMCs were largely sex-specific with only 15 age-related DMCs overlapping between both male and female control animals. Detailed age-related DMCs split by sex are available as supplementary data tables (Supplementary File 9-10).

**Table 4:** Age-related and deflected DMCs. Age-related DMCs have differential modification between 6 and 36 weeks, detected using the DSS R package; FDR<0.05. Deflected DMCs are age-related DMCs with a significant interaction between age and exposure, detected using the DSS R package; FDR<0.05.

| Age-related DMCs |  |  |
| --- | --- | --- |
| Direction | Female | Male |
| Hypermethylated | 143 | 89 |
| Hypomethylated | 265 | 201 |
| **Total** | **408** | **290** |
| Deflected DMCs |  |  |
| Direction | Female | Male |
| Hyper to hypo | 71 | 15 |
| Hypo to hyper | 29 | 3 |
| Hyper to hyper | 2 | 0 |
| Hypo to hypo | 0 | 0 |

Switch Premature aging

To reduce the number of comparisons and preserve statistical power during differential testing across multiple time points, only those cytosines with significant age-related changes were included in subsequent age:exposure interaction modeling. In the age:exposure interaction models, we identified 102 DMCs in female samples and 18 in male samples that had a significant interaction between age and exposure (Table 4, Supplementary Files 11-12). Of note, none of these “deflected DMCs” overlapped between the two sexes. Most of these DMCs were not previously identified by the cross-sectional analysis and map to 8 (female) and 2 (male) additional genes.

Next, we compared the direction of change for these “deflected DMCs” from 6 weeks to 36 weeks of age to determine if dieldrin led to premature epigenetic aging. Only 2 of these DMCs showed a pattern consistent with premature epigenetic aging. In this scenario, both DMCs were hypermodified at both 6 and 36 weeks by dieldrin, but the change in dieldrin animals was much smaller from 6 to 36 weeks, such that the β-value in dieldrin exposed animals at 6 weeks was “prematurely” high and more similar to the β-value in control animals at 36 weeks (Table 4). In contrast, most of these “deflected DMCs” show a “switching” pattern (i.e. inconsistent direction of change in age-related DNA modifications by exposure group) (Table 4). For example, a DMC that is hypermodified at 6 weeks in dieldrin exposed animals but hypomodified at 36 weeks, follows a switching pattern (hyper to hypo in Table 4).

**Figure 7:**
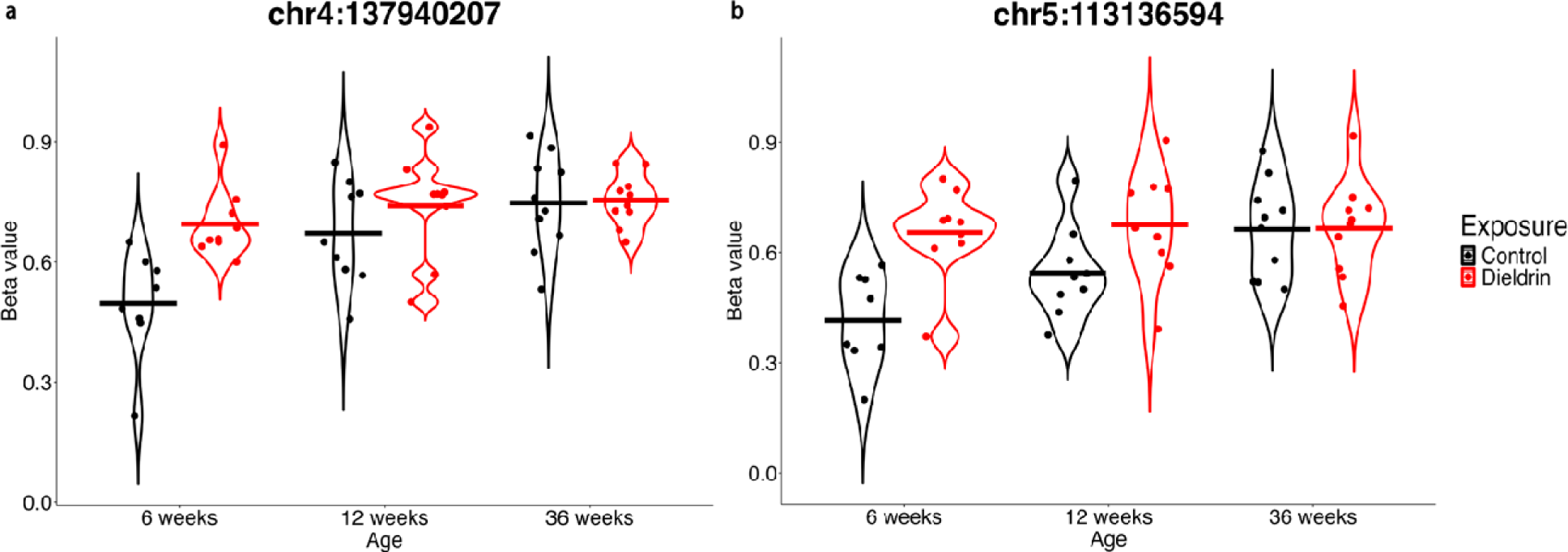
Raw beta values of the two premature aging DMCs. Violin plots display the raw beta values for control and dieldrin female samples at 6 weeks, 12 weeks, and 36 weeks for the deflected DMCs that show patterns consistent with premature epigenetic aging located at (a) chromosome 4, position 137940207 and (b) chromosome 5, position 113136594. Each point represents a single sample. The horizontal crossbars indicate the mean.

### Dieldrin is detectable in brains of F1 neonates

In addition to our epigenetic analysis, we tested whether dieldrin was present in the brains of F1 pups at birth. Previous data show that dieldrin is not detectable in the brain of F1 pups at 12 weeks of age in this exposure paradigm (Richardson *et al*., 2006). However, dieldrin can cross the placenta and the blood-brain barrier, so we expect dieldrin to be present in the brain during the exposure period. We collected brain tissue from F1 neonates (n=4 per group) from independent litters and measured dieldrin levels by mass spectrometry. As expected, dieldrin was detectable in dieldrin-exposed pups but not in vehicle-exposed pups, with a mean level of 621.9 ng dieldrin/g tissue in dieldrin-exposed offspring (Figure 8). Given previous estimates of the half-life of dieldrin in mouse at 1-10 days and approximately three days in mouse brain, our data that dieldrin is detectable in neonatal brain is consistent with previous data showing the dieldrin is not detectable by 12 weeks of age, which is 9 weeks after the end of the exposure period (Anonymous 1989,Hatcher *et al*., 2007,Richardson *et al*., 2006).

**Figure 8:**
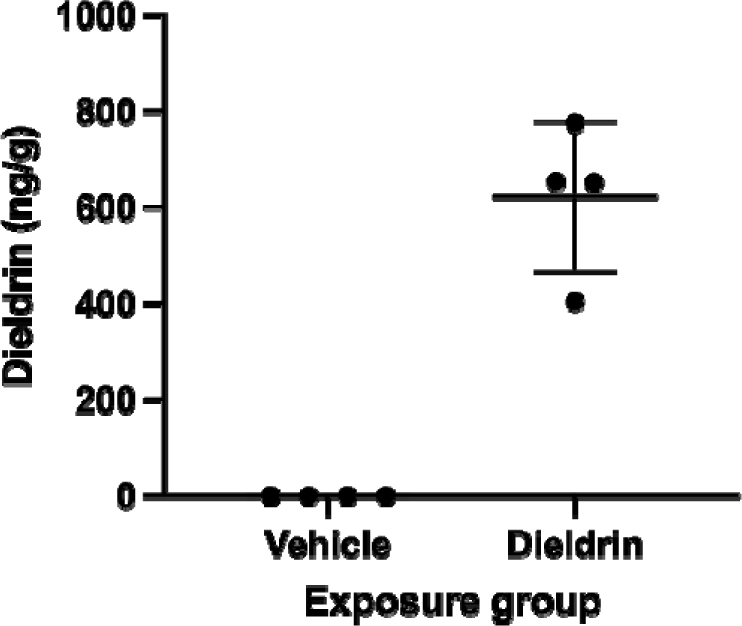
Dieldrin levels in F1 neonatal brain. Dieldrin levels in neonatal brain from vehicle- and dieldrin-exposed F1 animals. Data shown as mean ± SD.

## Discussion

### Dieldrin-induced differential modifications occur within genes with critical functions in neurodevelopment and in maintenance of neurological function

Our findings reinforce the concept of silent neurotoxicity, where the effects of developmental exposures are unmasked by challenges later in life, the cumulative effects of exposures over the lifespan, or the effects of aging (Cory-Slechta *et al*., 2005,Kraft *et al*., 2016). The epigenetic changes identified implicate potential mechanisms by which dieldrin primes the nigrostriatal system to have an exacerbated response to PD-related toxicity without observable changes in typical markers of nigrostriatal dysfunction and degeneration. The identification of DNA modifications in genes with critical roles in neurodevelopmental pathways and proper neurological function postnatally and into adulthood supports our hypothesis that developmental dieldrin exposure epigenetically regulates specific essential developmental programs and pathways with impacts that persist throughout the lifespan (Table 3, Figure 5,Figure 6). Here, we highlight specific differentially modified (DM) genes and pathways that may be important mediators of the persistent effect of dieldrin exposure (Figure 6, Table 3, Supplementary Files 6,7).

Two DM genes, *Lmx1b* and *Nr4a2* encode transcription factors involved in the development, maintenance, and survival of DA neurons, and in PD specifically (Arenas *et al*., 2015,Oliveira *et al*., 2017,Doucet-Beaupré *et al*., 2015,Lim *et al*., 2024,Gamit *et al*., 2023,Decressac *et al*., 2013,Jakaria *et al*., 2019,Al-Nusaif *et al*., 2022). Further, in our data, multiple components of fibroblast growth factor signaling (*Fgf8*, *Fgfr2,* and *Stat1*) are differentially modified. FGF signaling plays important roles in multiple aspects of neurodevelopment and neurological function in the developing and adult brain, including growth and patterning of the developing brain, neurogenesis, gliogenesis, axon outgrowth, myelination, tissue repair, synaptogenesis, synapse maturation, astrocyte-mediated synaptic pruning, and the maintenance of glia-neuron interactions (Scheltinga *et al*., 2013,Stevens *et al*., 2010,Klimaschewski and Claus, 2021,Dabrowski *et al*., 2015,Stevens *et al*., 2023). Supporting the idea that synaptogenesis pathways are affected by early-life dieldrin exposure, additional DM genes have known functions related to synaptogenesis, the maintenance and function of synapses, and synaptic plasticity, including *Ephb2*, *Nrxn1*, *Dlg2*, *Dlgap1*, *Camk2b*, *Prkca*, and *Prkce* (Kania and Klein, 2016,Sloniowski and Ethell, 2012,Assali *et al*., 2021,Liu *et al*., 2022,Südhof, 2023,Borghi *et al*., 2023,Kaizuka and Takumi, 2024,Sen *et al*., 2016,Rasmussen *et al*., 2017,Luderman *et al*., 2015,Bu *et al*., 2021).Together, these pathways coordinate the formation, structure, function, and plasticity of synapses across the lifespan, consistent with the idea that the integrity of synapses is critical for proper neurotransmission and that multiple PD-related mechanism converge on disruption of synaptic function (Alter *et al*., 2013,Brooker *et al*., 2024,Soukup *et al*., 2018).

Given the well-documented roles of these genes in these processes, dieldrin-induced changes in epigenetic regulation over time in these genes could contribute to altered susceptibility of the dopaminergic system. It is possible that the observed alterations in genomic and temporal patterns of DNA modifications in these genes change how gene expression is regulated longitudinally, with differing effects on function over time. These data do not yet explore how this epigenetic regulation affects expression of these genes over time or in response to a parkinsonian insult. Future studies will determine if epigenetic regulation of these loci affects gene expression and transcript usage over time at the RNA and protein levels, as almost all these highlighted genes have multiple promoters and/or transcripts. In addition, we can utilize our two-hit dieldrin/PFF model to elucidate mechanisms by which those epigenetic changes impact the regulation of expression of these genes in a PD-related model.

### Differentially modified genes play important roles in glial cell function and neuroinflammation

Importantly, many of the identified DM genes highlighted above have known expression and functions in glial cells and in neuroinflammatory processes (Table 3). For example, FGF signaling mediates glia-neuron interactions including those between oligodendrocytes and axons that are critical for myelination and between astrocytes and synapses for synaptic pruning and plasticity (Stevens *et al*., 2023,Klimaschewski and Claus, 2021). In addition to its role as a synaptic organizing protein, NRXN1 also regulates spine morphology, synaptic strength, and myelination of axons by mediating glia-neuron interactions with astrocytes, oligodendrocytes, and microglia (Liu *et al*., 2022). Along with its roles in neuron-neuron signaling, recent evidence shows potential pro-inflammatory roles for EPHB2 in multiple types of glial cells, possibly via activation by TNF-α/NF-_Κ_B (Darling and Lamb, 2019,Pozniak *et al*., 2014,Ernst *et al*., 2019,Yang, J. *et al*., 2018). NF-_Κ_B has been well-studied in PD and has emerged as a potential mediator in the effects of toxicant exposures on neuroinflammation in PD (Anderson *et al*., 2018). Together, these results add to the rapidly growing recognition of the multifaceted functions of astrocytes, microglia, oligodendrocytes, and other non-neuronal cells, and highlights the need for additional studies into the role of glial cells and neuroinflammation in the response to neurotoxicants and PD.

### Distinct sex-specific responses to exposures may underly sex-specific differences in disease

The reported epigenetic effects are largely sex-specific, adding to a growing body of evidence that sex differences in PD could be due to sex-specific responses to PD-related toxicants (Kochmanski *et al*., 2019,Gezer *et al*., 2020,Richardson *et al*., 2006,Adamson *et al*., 2022). Sex dimorphisms in PD incidence are well-documented, as is a male-specific relative vulnerability to PD-related toxicants (Gillies *et al*., 2014,Jurado-Coronel *et al*., 2018,Haaxma *et al*., 2007,Dluzen and McDermott, 2000,Cerri *et al*., 2019,Eeden *et al*., 2003,Georgiev *et al*., 2017,Adamson *et al*., 2022). Multiple mechanisms have been proposed as putative mediators of these sex-specific effects, including but not limited to sex differences in glial cell function and immune response (Jurado-Coronel *et al*., 2018,Gillies *et al*., 2014,Adamson *et al*., 2022). In line with this, our previous study identified sex-specific effects of dieldrin on neuroinflammatory gene expression and the data here expand on that with sex-specific effects on DNA modification in genes with critical functions in glia (Gezer *et al*., 2020). More generally, these data add to evidence of the sex-specific nature of epigenetic responses to developmental exposures, reiterating the importance of considering sex when investigating the epigenetic mechanisms driving disease development (Svoboda *et al*., 2022b,Hilz and Gore, 2022,McCabe *et al*., 2017).

### Dieldrin induces age- and location-specific changes in DNA modifications

In addition to the sex-specificity of dieldrin-induced differential modification, our data highlight the age- and location-specificity of DNA modifications (Figure 6). By using capture hybridization-sequencing, we were able to gain a more complete map of DNA modifications across modified genes identified in a previous study in our group; these data show that even within one gene, the direction of change is highly dependent on the specific location (Kochmanski *et al*., 2019). In addition, the inclusion of multiple time points demonstrates that the position and direction of differential modifications is highly dependent on age. In particular, the imprinted genes, *Grb10* and *Gnas,* provide clear examples of the sex-, age- and location-specificity of differential modifications (Figure 6). Overall, dieldrin-associated alterations in the direction and location of the epigenetic regulation of these loci over time may alter gene expression, promoter usage, or isoform expression of these genes differently at different ages. These findings underscore the complexity of epigenetic regulation and the importance of studying the effects of exposures longitudinally, as the epigenetic regulation of these genes changes over time.

### Dieldrin exposure does not induce large-scale deflection of long-term trajectories of age-related DNA modification patterns

Contrary to the environmental deflection hypothesis, our analysis identified very few DMCs with dieldrin-associated deflection of age-related trajectories (Table 4) (Kochmanski *et al*., 2017). While this suggests that dieldrin may not be causing large-scale deflection of age-related trajectories in DNA modification patterns, this is at odds with the age-related differences identified in our cross-sectional analysis and multiple issues limit such a broad interpretation. First, this could be due to statistical differences in how variance is calculated for these two tests within the *DSS* package, which would explain why we found few age-related changes identified in control animals. Second, the birth time point was not included in this analysis because 1) the birth time point was collected from a different cohort and 2) because neonatal brains are much smaller, these microdissections are quite different. We would expect to see more age-related changes from birth to 36 weeks, but these experimental issues complicate the interpretation of that comparison. Third, because we used a targeted method to look at a small fraction of the genome, we may have missed many age-related changes. Finally, this analysis would omit DMCs without age-related change in control animals but with age-related change in dieldrin animals. However, limiting the analysis here was critical to preserve statistical power. Overall, it is difficult to make broad conclusions from these data, and the potential interaction between developmental exposure and epigenetic aging warrants further investigation.

## Acknowledgements

The authors thank Marie Adams and the Van Andel Genomics Core for providing consultation, library preparation, and next-generation sequencing facilities and services (RRID:SCR_022913). We also thank Cassandra Johnny and the Michigan State University Mass Spectrometry and Metabolomics Core for providing mass spectrometry services. We also thank Daofeng Li from Washington University in St. Louis for creating a mouse ChromHMM track at our request for the WashU Epigenome Browser.

## Data availability

Preregistration: https://doi.org/10.17605/OSF.IO/3QXFV

Raw sequencing data: GEO Accession No. GSE264165 (embargoed until publication)

All code, processed data, and supporting data will be available as Supplementary Files in the Dryad Data Repository upon publication: https://doi.org/10.5061/dryad.ffbg79d30

## CRediT Author Statement

**Joseph Kochmanski:** Conceptualization, Methodology, Software, Formal Analysis, Visualization, Investigation, Data Curation, Writing - Original Draft; **Mahek Virani:** Software, Validation, Formal Analysis, Data Curation, Writing - Review & Editing, Visualization**; Nathan C. Kuhn:** Software, Formal Analysis, Investigation**; Sierra L Boyd:** Investigation; **Marie Adams:** Investigation, Methodology; **Katelyn Becker:** Investigation; **Alison I. Bernstein:** Conceptualization, Methodology, Data Curation, Writing - Review & Editing, Supervision, Project, Visualization, Administration, Funding Acquisition

