## Supplementary Tables and Figures for "Developmental origins of Parkinson’s disease risk: perinatal exposure to the organochlorine pesticide dieldrin leads to sex-specific DNA modifications in critical neurodevelopmental pathways in the mouse midbrain"

**Supplementary Table 1: GO Terms enriched in female animals by time point.** Results of ClueGO gene ontology enrichment analysis and significant GO terms are shown ($p < 0.05$). GO terms are grouped when they share >50% of their genes. "% Associated Genes" indicates the percentage of the input genes associated with the term compared with all the genes associated with the term. Genes present at all time points are indicated in bold.

| GO Term | GO Groups | % Associated Genes | Associated Genes Found |
|---|---|---|---|
| **Birth** | | | |
| cell fate specification | Group0 | 5.22 | [Apc2, Dbx1, Dhh, Pax2, Tbx2, Tenm4] |
| adult locomotory behavior | Group1 | 5.61 | [Chd7, Inpp5f, Kcnj10, Mapt, Nr4a2, Sptbn4] |
| negative regulation of actin filament polymerization | Group2 | 4.05 | [Cracd, Fhod3, Sptbn4] |
| regulation of neuronal synaptic plasticity | Group3 | 6.67 | [Camk2b, **Ephb2**, Kcnj10, Ppfia3, Syap1] |
| filopodium assembly | Group4 | 6.17 | [**Ephb2**, **Myo3b**, Palm, **Ppp1r16b**, **Rhoq**] |
| regulation of filopodium assembly | Group4 | 8.06 | [**Ephb2**, **Myo3b**, Palm, **Ppp1r16b**, **Rhoq**] |
| negative regulation of neuron projection regeneration | Group5 | 16.67 | [Inpp5f, Rgma, **Xylt1**] |
| negative regulation of axon regeneration | Group5 | 18.75 | [Inpp5f, Rgma, **Xylt1**] |
| central nervous system neuron development | Group6 | 5.56 | [**Ephb2**, **Fgfr2**, Lmx1b, Nr4a2, **Prkca**, Sptbn4] |
| central nervous system neuron axonogenesis | Group6 | 8.89 | [**Ephb2**, Nr4a2, **Prkca**, Sptbn4] |
| ear morphogenesis | Group7 | 5.41 | [Chd7, **Ephb2**, **Fgfr2**, Hoxa2, **Myo3b**, Pax2, **Ptk7**, Tbx2] |
| inner ear morphogenesis | Group7 | 5.69 | [Chd7, **Ephb2**, **Fgfr2**, **Myo3b**, Pax2, **Ptk7**, Tbx2] |
| cochlea morphogenesis | Group7 | 14.29 | [**Myo3b**, Pax2, **Ptk7**, Tbx2] |
| **6 weeks** | | | |
| macrophage activation involved in immune response | Group00 | 14.29 | [Prkce, Sbno2, Syk] |
| cardiac myofibril assembly | Group01 | 13.64 | [Fhod3, Nrap, Srf] |
| coronary vasculature development | Group02 | 4 | [**Fgfr2**, **Ptk7**, Srf] |
| structural constituent of postsynapse | Group03 | 13.64 | [Camk2b, Dlg2, Dlgap1] |
| ear morphogenesis | Group04 | 4.73 | [**Ephb2**, Fgf8, **Fgfr2**, Hoxa2, **Myo3b**, **Ptk7**, Tshz1] |
| negative regulation of glucose import | Group05 | 13.64 | [Grb10, **Prkca**, **Rhoq**] |
| regulation of neuronal synaptic plasticity | Group06 | 4 | [Camk2b, **Ephb2**, Kcnj10] |
| regulation of mesenchymal cell proliferation | Group07 | 9.3 | [**Fgfr2**, Foxp1, Lmna, Stat1] |
| positive regulation of smooth muscle cell proliferation | Group07 | 5.17 | [Cdh13, Elane, **Fgfr2**, Foxp1, **Prkca**, Stat1] |
| embryonic skeletal system development | Group08 | 4.7 | [Fgf8, **Fgfr2**, Gnas, Hoxa2, Hoxa3, Hoxa4, **Xylt1**] |
| embryonic skeletal system morphogenesis | Group08 | 5.5 | [Fgf8, **Fgfr2**, Gnas, Hoxa2, Hoxa3, Hoxa4] |
| filopodium assembly | Group09 | 6.17 | [**Ephb2**, **Myo3b**, **Ppp1r16b**, **Rhoq**, Srf] |
| regulation of filopodium assembly | Group09 | 8.06 | [**Ephb2**, **Myo3b**, **Ppp1r16b**, **Rhoq**, Srf] |
| **12 weeks** | | | |
| positive regulation of smooth muscle cell proliferation | Group00 | 5.17 | [Cdh13, **Fgfr2**, Foxp1, Il13, Mdm2, **Prkca**] |
| ceramide biosynthetic process | Group01 | 4.23 | [Gal3st1, Smpd5, St3gal2] |
| activation of GTPase activity | Group02 | 4.46 | [Agap1, Apc2, Arhgap45, Sipa1l3, Tbc1d4] |
| cellular response to alcohol | Group03 | 4.46 | [Dnmt3a, Gnas, Gramd1c, Lrp8, Prkce] |
| protein localization to cell junction | Group04 | 4.72 | [Actn4, Dlg2, Dlgap1, Heg1, Nectin1, Tjp3] |
| central nervous system neuron development | Group05 | 4.63 | [**Ephb2**, **Fgfr2**, Lmx1b, Nr4a2, **Prkca**] |
| negative regulation of carbohydrate metabolic process | Group06 | 4.05 | [Cbfa2t3, Grb10, Midn] |
| filopodium assembly | Group07 | 6.17 | [**Ephb2**, **Myo3b**, Palm, **Ppp1r16b**, **Rhoq**] |
| regulation of filopodium assembly | Group07 | 8.06 | [**Ephb2**, **Myo3b**, Palm, **Ppp1r16b**, **Rhoq**] |

| | | | |
|---|---|---|---|
| negative regulation of glucose transmembrane transport | Group08 | 11.11 | [Grb10, **Prkca**, **Rhoq**] |
| negative regulation of glucose import | Group08 | 13.64 | [Grb10, **Prkca**, **Rhoq**] |
| negative regulation of neuron projection regeneration | Group09 | 16.67 | [Inpp5f, Rgma, **Xylt1**] |
| negative regulation of axon regeneration | Group09 | 18.75 | [Inpp5f, Rgma, **Xylt1**] |
| ear morphogenesis | Group10 | 4.73 | [**Ephb2**, **Fgfr2**, **Myo3b**, **Ptk7**, Tbx2, Tshz1, Ush1g] |
| inner ear morphogenesis | Group10 | 4.88 | [**Ephb2**, **Fgfr2**, **Myo3b**, **Ptk7**, Tbx2, Ush1g] |
| cochlea morphogenesis | Group10 | 10.71 | [**Myo3b**, **Ptk7**, Tbx2] |
| striated muscle cell proliferation | Group11 | 6.1 | [**Fgfr2**, Foxp1, Rxra, Tbx2, Tenm4] |
| heart growth | Group11 | 4.92 | [**Fgfr2**, Foxp1, Heg1, Rxra, Tbx2, Tenm4] |
| cardiac muscle tissue growth | Group11 | 5.41 | [**Fgfr2**, Foxp1, Heg1, Rxra, Tbx2, Tenm4] |
| coronary vasculature development | Group11 | 4 | [**Fgfr2**, **Ptk7**, Rxra] |
| cardiac muscle cell proliferation | Group11 | 8.06 | [**Fgfr2**, Foxp1, Rxra, Tbx2, Tenm4] |
| ventricular cardiac muscle tissue development | Group11 | 4.23 | [**Fgfr2**, Heg1, Rxra] |
| **36 weeks** | | | |
| ceramide biosynthetic process | Group0 | 4.23 | [Cyp4f39, Smpd5, St3gal2] |
| protein O-linked glycosylation | Group1 | 4.41 | [Galnt18, Galnt2, Pomgnt2] |
| platelet aggregation | Group2 | 4.84 | [Dmtn, Gnas, **Prkca**] |
| filopodium assembly | Group3 | 6.17 | [Dmtn, **Ephb2**, **Myo3b**, **Ppp1r16b**, **Rhoq**] |
| regulation of filopodium assembly | Group3 | 8.06 | [Dmtn, **Ephb2**, **Myo3b**, **Ppp1r16b**, **Rhoq**] |
| negative regulation of glucose transmembrane transport | Group4 | 11.11 | [Grb10, **Prkca**, **Rhoq**] |
| regulation of glucose import | Group4 | 4.29 | [Grb10, **Prkca**, **Rhoq**] |
| negative regulation of glucose import | Group4 | 13.64 | [Grb10, **Prkca**, **Rhoq**] |
| protein methyltransferase activity | Group5 | 4.21 | [Pcmtd1, Prdm16, Setd1b, Smyd3] |
| protein-lysine N-methyltransferase activity | Group5 | 5 | [Prdm16, Setd1b, Smyd3] |
| histone methyltransferase activity | Group5 | 5 | [Prdm16, Setd1b, Smyd3] |
| histone-lysine N-methyltransferase activity | Group5 | 6.82 | [Prdm16, Setd1b, Smyd3] |
| mesenchymal cell proliferation | Group6 | 5.17 | [**Fgfr2**, Foxp1, Lmna] |
| regulation of mesenchymal cell proliferation | Group6 | 6.98 | [**Fgfr2**, Foxp1, Lmna] |
| positive regulation of smooth muscle cell proliferation | Group6 | 4.31 | [Cdh13, **Fgfr2**, Foxp1, Map3k5, **Prkca**] |
| cardiac muscle cell proliferation | Group6 | 4.84 | [**Fgfr2**, Foxp1, Tenm4] |
| blood vessel remodeling | Group7 | 6.67 | [Atg5, Chd7, Fgf8, Hoxa3] |
| cranial skeletal system development | Group7 | 4.71 | [Fgf8, **Fgfr2**, Gnas, Hoxa2] |
| positive regulation of stem cell proliferation | Group7 | 4.62 | [Fgf8, **Fgfr2**, Hoxa3] |
| cranial nerve development | Group7 | 5 | [Chd7, **Ephb2**, Hoxa3] |
| hindlimb morphogenesis | Group7 | 8.33 | [Aff3, Chd7, Fgf8, Gnas] |
| embryonic cranial skeleton morphogenesis | Group7 | 7.27 | [Fgf8, **Fgfr2**, Gnas, Hoxa2] |
| embryonic hindlimb morphogenesis | Group7 | 10.26 | [Aff3, Chd7, Fgf8, Gnas] |
| motor neuron axon guidance | Group8 | 7.89 | [Fgf8, Foxp1, Hoxa2] |
| blood vessel remodeling | Group8 | 6.67 | [Atg5, Chd7, Fgf8, Hoxa3] |
| cranial skeletal system development | Group8 | 4.71 | [Fgf8, **Fgfr2**, Gnas, Hoxa2] |
| positive regulation of stem cell proliferation | Group8 | 4.62 | [Fgf8, **Fgfr2**, Hoxa3] |
| cardiac septum development | Group8 | 4.03 | [Chd7, Fgf8, **Fgfr2**, Heg1, **Ptk7**] |
| cellular response to retinoic acid | Group8 | 4.48 | [**Fgfr2**, Hoxa2, **Ptk7**] |
| hindlimb morphogenesis | Group8 | 8.33 | [Aff3, Chd7, Fgf8, Gnas] |
| ear morphogenesis | Group8 | 4.73 | [Chd7, **Ephb2**, Fgf8, **Fgfr2**, Hoxa2, **Myo3b**, **Ptk7**] |

| | | | |
|---|---|---|---|
| inner ear morphogenesis | Group8 | 4.88 | [Chd7, **Ephb2**, Fgf8, **Fgfr2**, **Myo3b**, **Ptk7**] |
| embryonic skeletal system development | Group8 | 4.03 | [Fgf8, **Fgfr2**, Gnas, Hoxa2, Hoxa3, **Xylt1**] |
| embryonic skeletal system morphogenesis | Group8 | 4.59 | [Fgf8, **Fgfr2**, Gnas, Hoxa2, Hoxa3] |
| embryonic cranial skeleton morphogenesis | Group8 | 7.27 | [Fgf8, **Fgfr2**, Gnas, Hoxa2] |
| ventricular cardiac muscle tissue development | Group8 | 4.23 | [Chd7, **Fgfr2**, Heg1] |
| embryonic hindlimb morphogenesis | Group8 | 10.26 | [Aff3, Chd7, Fgf8, Gnas] |
| central nervous system neuron development | Group8 | 4.63 | [**Ephb2**, Fgf8, **Fgfr2**, Nr4a2, **Prkca**] |
| ventricular cardiac muscle tissue morphogenesis | Group8 | 4.92 | [Chd7, **Fgfr2**, Heg1] |
| central nervous system neuron axonogenesis | Group8 | 6.67 | [**Ephb2**, Nr4a2, **Prkca**] |

**Supplementary Table 2: GO Terms enriched in male animals by time point**. Results of ClueGO gene ontology enrichment analysis and significant GO terms are shown ($p < 0.05$). GO terms are grouped when they share >50% of their genes. "% Associated Genes" indicates the percentage of the input genes associated with the term compared with all the genes associated with the term. Genes present at all time points are indicated in bold.

| Term | GO Groups | % Associated Genes | Associated Genes Found |
|---|---|---|---|
| **Birth** | | | |
| macrophage activation involved in immune response | Group00 | 14.29 | [Cx3cr1, **Prkce**, Sbno2] |
| ear morphogenesis | Group01 | 4.73 | [**Ephb2**, Fgf8, **Fgfr2**, Hoxa2, **Myo3b**, **Ptk7**, Tshz1] |
| positive thymic T cell selection | Group02 | 15.79 | [Cd3d, Srf, Stk11] |
| negative regulation of carbohydrate metabolic process | Group03 | 4.05 | [Cbfa2t3, Grb10, Midn] |
| glycerolipid catabolic process | Group04 | 4.17 | [Inpp5f, Mgll, Pla2g7] |
| synapse maturation | Group05 | 12.12 | [Camk2b, Cx3cr1, Nrxn1, Palm] |
| lung morphogenesis | Group06 | 4.17 | [Fgf8, **Fgfr2**, Srf] |
| regulation of neurotransmitter receptor activity | Group07 | 4.17 | [Dlgap1, **Ephb2**, Nrxn1] |
| protein localization to cell junction | Group08 | 5.51 | [Actn4, Dlg2, Dlgap1, Heg1, Nectin1, Nrxn1, Tjp3] |
| osteoclast differentiation | Group09 | 4.8 | [Efna2, Foxp1, Fstl3, **Gnas**, Sbno2, Sh3pxd2a] |
| filopodium assembly | Group10 | 8.64 | [Dmtn, **Ephb2**, **Myo3b**, Nrxn1, Palm, **Ppp1r16b**, Srf] |
| regulation of filopodium assembly | Group10 | 11.29 | [Dmtn, **Ephb2**, **Myo3b**, Nrxn1, Palm, **Ppp1r16b**, Srf] |
| negative regulation of neuron projection regeneration | Group11 | 16.67 | [Inpp5f, Rgma, Xylt1] |
| negative regulation of axon regeneration | Group11 | 18.75 | [Inpp5f, Rgma, Xylt1] |
| peptidyl-threonine modification | Group12 | 5.97 | [Dmtn, Galnt2, **Myo3b**, Ppp2r5d, Prkag2, **Prkca**, Stk11, Ttbk1] |
| peptidyl-threonine phosphorylation | Group12 | 5.6 | [Dmtn, **Myo3b**, Ppp2r5d, Prkag2, **Prkca**, Stk11, Ttbk1] |
| regulation of cell morphogenesis involved in differentiation | Group13 | 4.76 | [Actn4, Camk2b, Dmtn, **Ephb2**, Lrp8, Meltf] |
| negative regulation of cell morphogenesis involved in differentiation | Group13 | 20 | [Actn4, Dmtn, Meltf] |
| negative regulation of substrate adhesion-dependent cell spreading | Group13 | 21.43 | [Actn4, Dmtn, Meltf] |
| **6 weeks** | | | |
| dopamine receptor signaling pathway | Group00 | 7.32 | [Gnao1, **Gnas**, Palm] |
| cellular response to heat | Group01 | 4.92 | [Ano1, Arpp21, Xylt1] |
| collagen catabolic process | Group02 | 6 | [Mmp28, Prtn3, Retreg1] |
| lipopolysaccharide-mediated signaling pathway | Group03 | 4.62 | [**Prkca**, **Prkce**, Stat1] |
| phospholipid dephosphorylation | Group04 | 6.82 | [Inpp5a, Inpp5f, Plppr3] |
| protein phosphatase regulator activity | Group05 | 4.35 | [Elfn2, **Ppp1r16b**, Ppp2r5c, Ppp2r5d] |
| cellular component maintenance | Group06 | 4.4 | [Dlg2, Dlgap1, Prtn3, Tanc1] |
| cell junction maintenance | Group06 | 4.92 | [Dlg2, Dlgap1, Prtn3] |
| mesenchymal cell proliferation | Group07 | 5.17 | [**Fgfr2**, Lmna, Stat1] |
| regulation of mesenchymal cell proliferation | Group07 | 6.98 | [**Fgfr2**, Lmna, Stat1] |
| cellular response to retinoic acid | Group08 | 4.48 | [**Fgfr2**, Hoxa2, **Ptk7**] |
| embryonic cranial skeleton morphogenesis | Group08 | 5.45 | [**Fgfr2**, **Gnas**, Hoxa2] |
| peptidyl-threonine modification | Group09 | 4.48 | [Galnt2, **Myo3b**, Ppp2r5d, Prkag2, **Prkca**, Tbk1] |
| peptidyl-threonine phosphorylation | Group09 | 4 | [**Myo3b**, Ppp2r5d, Prkag2, **Prkca**, Tbk1] |
| filopodium assembly | Group10 | 6.17 | [**Ephb2**, **Myo3b**, Palm, **Ppp1r16b**, Rhoq] |

| | | | |
|---|---|---|---|
| regulation of filopodium assembly | Group10 | 8.06 | [**Ephb2**, **Myo3b**, Palm, **Ppp1r16b**, Rhoq] |
| positive regulation of filopodium assembly | Group10 | 7.5 | [**Myo3b**, Palm, Rhoq] |
| negative regulation of cellular response to insulin stimulus | Group11 | 6.52 | [Grb10, **Prkca**, Tns2] |
| negative regulation of glucose transmembrane transport | Group11 | 11.11 | [Grb10, **Prkca**, Rhoq] |
| negative regulation of insulin receptor signaling pathway | Group11 | 7.32 | [Grb10, **Prkca**, Tns2] |
| regulation of glucose import | Group11 | 4.29 | [Grb10, **Prkca**, Rhoq] |
| negative regulation of glucose import | Group11 | 13.64 | [Grb10, **Prkca**, Rhoq] |
| **12 weeks** | | | |
| lymph vessel development | Group0 | 9.09 | [Heg1, Ptpn14, Syk] |
| macrophage activation involved in immune response | Group1 | 14.29 | [**Prkce**, Sbno2, Syk] |
| negative regulation of carbohydrate metabolic process | Group2 | 4.05 | [Cbfa2t3, Grb10, Midn] |
| negative regulation of cellular carbohydrate metabolic process | Group2 | 4.41 | [Cbfa2t3, Grb10, Midn] |
| filopodium assembly | Group3 | 4.94 | [**Ephb2**, **Myo3b**, Palm, **Ppp1r16b**] |
| regulation of filopodium assembly | Group3 | 6.45 | [**Ephb2**, **Myo3b**, Palm, **Ppp1r16b**] |
| mesenchymal cell proliferation | Group4 | 5.17 | [**Fgfr2**, Foxp1, Lmna] |
| regulation of mesenchymal cell proliferation | Group4 | 6.98 | [**Fgfr2**, Foxp1, Lmna] |
| cardiac muscle cell proliferation | Group4 | 4.84 | [**Fgfr2**, Foxp1, Tenm4] |
| regulation of platelet activation | Group5 | 5.66 | [Gp1bb, **Prkca**, Syk] |
| platelet aggregation | Group5 | 4.84 | [**Gnas**, **Prkca**, Syk] |
| positive regulation of cold-induced thermogenesis | Group5 | 5 | [**Gnas**, Grb10, Pemt, Prdm16, Syk] |
| regulation of response to reactive oxygen species | Group6 | 6.82 | [Foxo3, Foxp1, Stk26] |
| cell death in response to hydrogen peroxide | Group6 | 7.32 | [Foxo3, Foxp1, Stk26] |
| regulation of hydrogen peroxide-induced cell death | Group6 | 8.11 | [Foxo3, Foxp1, Stk26] |
| motor neuron axon guidance | Group7 | 7.89 | [Fgf8, Foxp1, Hoxa2] |
| cranial skeletal system development | Group7 | 4.71 | [Fgf8, **Fgfr2**, **Gnas**, Hoxa2] |
| animal organ formation | Group7 | 5.06 | [Fgf8, **Fgfr2**, Hoxa3, Pax2] |
| cochlea development | Group7 | 4.84 | [**Myo3b**, Pax2, **Ptk7**] |
| specification of animal organ identity | Group7 | 6.98 | [Fgf8, **Fgfr2**, Pax2] |
| positive regulation of stem cell proliferation | Group7 | 4.62 | [Fgf8, **Fgfr2**, Hoxa3] |
| cranial nerve morphogenesis | Group7 | 9.68 | [**Ephb2**, Hoxa3, Pax2] |
| cellular response to retinoic acid | Group7 | 5.97 | [**Fgfr2**, Hoxa2, Pax2, **Ptk7**] |
| cranial nerve development | Group7 | 5 | [**Ephb2**, Hoxa3, Pax2] |
| hindlimb morphogenesis | Group7 | 6.25 | [Aff3, Fgf8, **Gnas**] |
| ear morphogenesis | Group7 | 4.73 | [**Ephb2**, Fgf8, **Fgfr2**, Hoxa2, **Myo3b**, Pax2, **Ptk7**] |
| inner ear morphogenesis | Group7 | 4.88 | [**Ephb2**, Fgf8, **Fgfr2**, **Myo3b**, Pax2, **Ptk7**] |
| brain morphogenesis | Group7 | 6.67 | [Fgf8, Foxo3, Pax2] |
| embryonic skeletal system morphogenesis | Group7 | 4.59 | [Fgf8, **Fgfr2**, **Gnas**, Hoxa2, Hoxa3] |
| cochlea morphogenesis | Group7 | 10.71 | [**Myo3b**, Pax2, **Ptk7**] |
| embryonic cranial skeleton morphogenesis | Group7 | 7.27 | [Fgf8, **Fgfr2**, **Gnas**, Hoxa2] |
| embryonic hindlimb morphogenesis | Group7 | 7.69 | [Aff3, Fgf8, **Gnas**] |
| **36 weeks** | | | |
| regulation of alternative mRNA splicing, via spliceosome | Group00 | 4.48 | [Ptbp1, Rbfox1, Rbfox3] |
| platelet aggregation | Group01 | 4.84 | [**Gnas**, **Prkca**, Syk] |
| macrophage activation involved in immune response | Group02 | 14.29 | [**Prkce**, Sbno2, Syk] |
| establishment of epithelial cell polarity | Group03 | 7.5 | [Frmd4b, **Ptk7**, Sipa1l3] |
| chemosensory behavior | Group04 | 12 | [Chd7, Lmx1b, **Prkce**] |

| | | | |
|---|---|---|---|
| tumor necrosis factor-mediated signaling pathway | Group05 | 4.49 | [Actn4, Foxo3, Stat1, Syk] |
| neuron maturation | Group06 | 4.76 | [Dlg2, Epha8, Nr4a2] |
| positive regulation of calcium-mediated signaling | Group07 | 6.82 | [Cdh13, Ptbp1, Syk] |
| neuromuscular process controlling balance | Group08 | 4.41 | [Camk2b, Rbfox1, Ush1g] |
| regulation of filopodium assembly | Group09 | 4.84 | [**Ephb2**, **Myo3b**, **Ppp1r16b**] |
| syncytium formation | Group10 | 5.26 | [Sbno2, Sh3pxd2a, Stat1, Tanc1] |
| syncytium formation by plasma membrane fusion | Group10 | 5.41 | [Sbno2, Sh3pxd2a, Stat1, Tanc1] |
| positive regulation of synapse assembly | Group11 | 4.94 | [**Ephb2**, Flrt1, Lrrc4b, **Prkca**] |
| dopaminergic neuron differentiation | Group11 | 6.52 | [Lmx1b, Nr4a2, Pax2] |
| central nervous system neuron development | Group11 | 4.63 | [**Ephb2**, **Fgfr2**, Lmx1b, Nr4a2, **Prkca**] |
| central nervous system neuron axonogenesis | Group11 | 6.67 | [**Ephb2**, Nr4a2, **Prkca**] |
| regulation of neuronal synaptic plasticity | Group12 | 4 | [Camk2b, **Ephb2**, Kcnj10] |
| regulation of long-term neuronal synaptic plasticity | Group12 | 8.11 | [Camk2b, **Ephb2**, Kcnj10] |
| positive regulation of dendritic spine development | Group12 | 4.69 | [Camk2b, **Ephb2**, Itsn1] |
| positive regulation of dendrite morphogenesis | Group12 | 4.84 | [Camk2b, Cul7, **Ephb2**] |
| cochlea development | Group13 | 6.45 | [**Myo3b**, Pax2, **Ptk7**, Tbx2] |
| cellular response to retinoic acid | Group13 | 4.48 | [**Fgfr2**, Pax2, **Ptk7**] |
| cranial nerve development | Group13 | 5 | [Chd7, **Ephb2**, Pax2] |
| ear morphogenesis | Group13 | 5.41 | [Chd7, **Ephb2**, **Fgfr2**, **Myo3b**, Pax2, **Ptk7**, Tbx2, Ush1g] |
| inner ear morphogenesis | Group13 | 6.5 | [Chd7, **Ephb2**, **Fgfr2**, **Myo3b**, Pax2, **Ptk7**, Tbx2, Ush1g] |
| cochlea morphogenesis | Group13 | 14.29 | [**Myo3b**, Pax2, **Ptk7**, Tbx2] |
| mesenchymal cell proliferation | Group14 | 8.62 | [**Fgfr2**, Foxp1, Lmna, Stat1, Tbx2] |
| striated muscle cell proliferation | Group14 | 4.88 | [**Fgfr2**, Foxp1, Tbx2, Tenm4] |
| regulation of mesenchymal cell proliferation | Group14 | 9.3 | [**Fgfr2**, Foxp1, Lmna, Stat1] |
| positive regulation of mesenchymal cell proliferation | Group14 | 8.57 | [**Fgfr2**, Foxp1, Stat1] |
| positive regulation of smooth muscle cell proliferation | Group14 | 4.31 | [Cdh13, **Fgfr2**, Foxp1, **Prkca**, Stat1] |
| lipopolysaccharide-mediated signaling pathway | Group14 | 4.62 | [**Prkca**, **Prkce**, Stat1] |
| cardiac muscle cell proliferation | Group14 | 6.45 | [**Fgfr2**, Foxp1, Tbx2, Tenm4] |
| regulation of cardiac muscle cell proliferation | Group14 | 6.12 | [**Fgfr2**, Foxp1, Tbx2] |

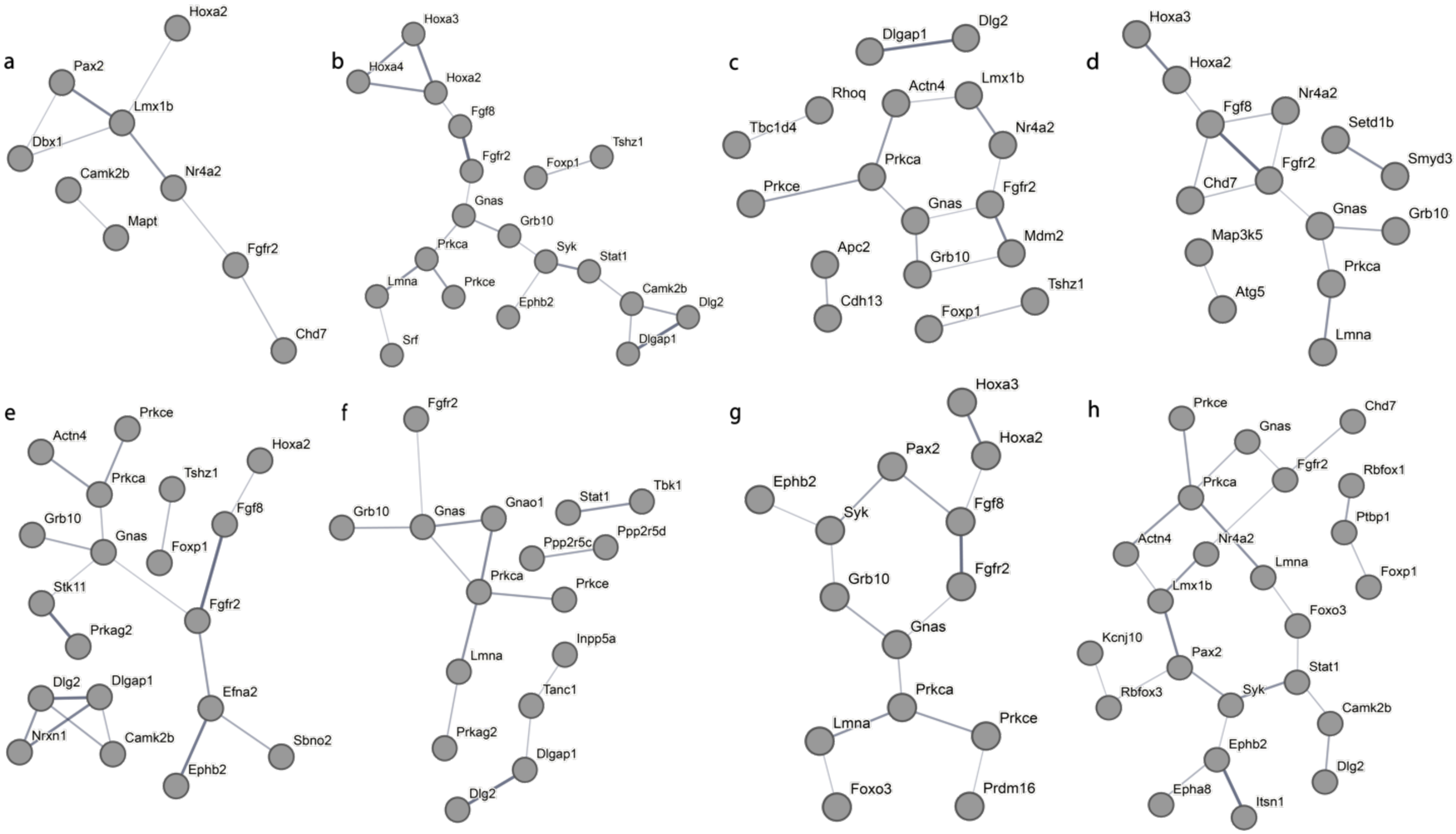


**Supplementary Figure 1: STRING protein-protein interaction networks between genes found in enriched GO terms.** Interaction networks for female (a-d) and male (e-h) at birth (a,e), 6 weeks (b,f), 12 weeks (c,g), and 36 weeks (d,h). STRING networks display the interactions with at least a 0.4 interaction score and omit any disconnected nodes. Each node represents all proteins produced by a single, protein coding gene locus. Edges represent protein-protein associations, but not necessarily physical interactions. Thickness of the lines indicates the strength of data support.
